# Deciphering selection patterns of somatic copy-number events

**DOI:** 10.64898/2026.03.01.708809

**Authors:** Tom L. Kaufmann, Adam Streck, Florian Markowetz, Peter Van Loo, Roland F. Schwarz

## Abstract

Somatic copy-number alterations (SCNAs) are pervasive across cancers, arising through diverse mutational mechanisms and subsequently shaped by selection according to fitness advantages [1–4]. These alterations can drive tumourigenesis via gene-dosage changes in oncogenes and tumour suppressors [5, 6]. However, observed SCNA profiles do not uniquely define the genomic events that led to them, which is a major obstacle to understanding the selective forces shaping cancer genomes.

Here we present SPICE, Selection Patterns In somatic Copy-number Events, an event-level framework that infers discrete copy-number events from allele-specific profiles and models focal selection from first principles, incorporating uniform breakpoint formation and locus-specific selective pressures. Applied to 5,966 samples from the TCGA dataset, SPICE quantifies the full spectrum of SCNA events, recapitulates known mutational processes, and tracks systematic shifts in event types before and after whole-genome duplication (WGD). Next, SPICE employs a generative selection model to identify the location of oncogenes and tumour-suppressors. Unlike previous approaches that detected peaks in aggregated copy-number signals, our model operates directly on inferred events and treats genome-wide breakpoint formation as a neutral reference against which locus-specific selection is detected. This analysis reveals 460 loci under selection which are highly enriched for known oncogenes and tumour suppressors, recapitulating most previously reported sites and uncovering many novel regions, and simultaneously showing that most internal copy-number events are not subject to focal selection.

These results establish a unified framework that deconvolves copy-number profiles into their underlying evolutionary events and greatly expand the catalogue of loci implicated in cancer development.

**Graphical abstract:** 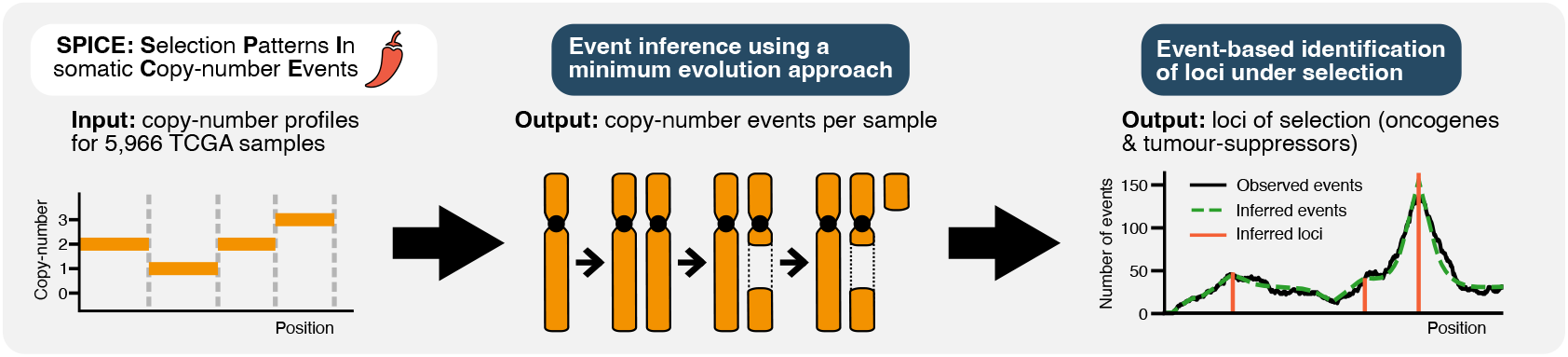

## Introduction

Cancer genomes harbour numerous somatic copy-number alterations (SCNAs), a hallmark of chromosomal instability that includes frequent whole-chromosome and whole-arm copy-number alterations, as well as segmental copy-number changes [1–3].

SCNAs arise via diverse mechanisms—including chromosome segregation errors [7], homologous recombination deficiency [8, 9], structural variants (SV) and complex genomic rearrangements [10, 11], whole-genome duplication [12] and chromothripsis [13]. Recent pan-cancer signature analyses have linked distinct SCNA patterns to some of these processes [14–16].

While SCNAs exhibit strong heterogeneity both between and within patients [17, 18], they also show clear signs of selective pressures during tumour evolution—such as parallel and convergent evolution [19, 20], tissue-specific patterns, and recurrent temporal sequences [21]. SCNAs can drive tumourigenesis through gene dosage effects by gaining oncogenes and losing tumour suppressor genes [5, 6, 22]. SCNAs shape the karyotype of a cell, that is, the number and structure of its chromosomes [23]. They are typically represented and analysed through copy-number profiles; however, these profiles do not map uniquely to a single karyotype. Each profile can correspond to multiple possible karyotypes, making the inferred copy-number features—such as segment lengths, event counts, and event types—potentially ambiguous (Supplementary Figure 1a). Horizontal dependencies between neighbouring segments further obscure focal targets, complicating the separation of mutations and therefore the detection of loci under selection.

Several methods have sought to disentangle these effects, most notably GISTIC, and BISCUT [6, 22, 24, 25]. These methods are limited either by reliance on coarse, thresholded recurrence summaries that cannot resolve complex overlapping events, or by restriction to events anchored at chromosome ends.

Here, we introduce **SPICE**, **S**election **P**atterns **I**n somatic **C**opy-number **E**vents, a framework that infers both discrete copy-number events and loci under selection from allelespecific profiles. SPICE infers event reconstructions via a reference-guided minimum-evolution procedure that explicitly accounts for whole-genome duplications. Across 5,966 samples from The Cancer Genome Atlas (TCGA), SPICE quantifies the full spectrum of SCNA events, recapitulates known mutational processes, and tracks systematic shifts in event types before and after whole-genome duplication (WGD). Building on this, SPICE employs a generative selection model that assumes uniform breakpoint formation and identifies loci where oncogenes or tumour suppressors distort event distributions, producing characteristic triangular shapes around loci under selection. This analysis reveals 460 loci under selection enriched for known and putative cancer genes, recovers the majority of previously reported calls, and highlights hundreds of novel regions. Notably, SPICE shows that most internal SCNAs are not subject to focal selection, yet exposes clear patterns of selection that depend on locus and event size, providing a unified view of mutational processes and selective forces shaping cancer genomes.

## Results

### Inferring copy-number events for 5,966 copynumber profiles

We inferred events from 5,966 high-quality samples spanning 33 cancer types from The Cancer Genome Atlas (TCGA; see Methods for a detailed description). To prepare the data for event inference, allele-specific copy-number profiles were obtained from the ASCAT v3 repository [26], preprocessed and filtered using CNSistent [27] and our custom pipeline (Methods), which included the removal of recurrent technical artefacts (Methods). We then split the 5,966 copy number profiles into allele-specific copy number profiles for individual chromosomes (46 per sample), excluded copy-number-neutral chromosomes, and obtained a final set of 157,214 allele-specific chromosome copy number profiles, each processed independently. Please note that subsequently, these allele-specific chromosomes are simply referred to as chromosomes.

In our framework, a copy-number event refers to a single gain or loss that alters the copy-number of a continuous chromosomal segment by +1 or -1. These events can vary in scale, from small segments spanning a few kilobases to entire chromosome arms or full chromosomes. Inferring such events is inherently challenging, as each observed profile can correspond to multiple possible karyotypes and evolutionary histories. To tackle this, we developed a new approach that leverages our minimum-evolution framework [28, 29], which parsimoniously reconstructs the fewest alterations consistent with the observed profile, enabling precise event-level resolution of copy-number changes (Figure 1a). Whole-genome duplication (WGD) events, which double the genomic content [30, 31], were explicitly modelled, with events classified as occurring either before or after the duplication. Samples showing evidence of multiple WGDs [32] were excluded. Next, each chromosome copy-number profile was encoded as a bipartite graph where nodes correspond to breakpoints and vertices correspond to gain or loss events connecting two break-points. Each possible minimum-evolution solution then corresponded to a valid way of connecting the nodes in this bipartite graph (Methods).

**Figure 1.**
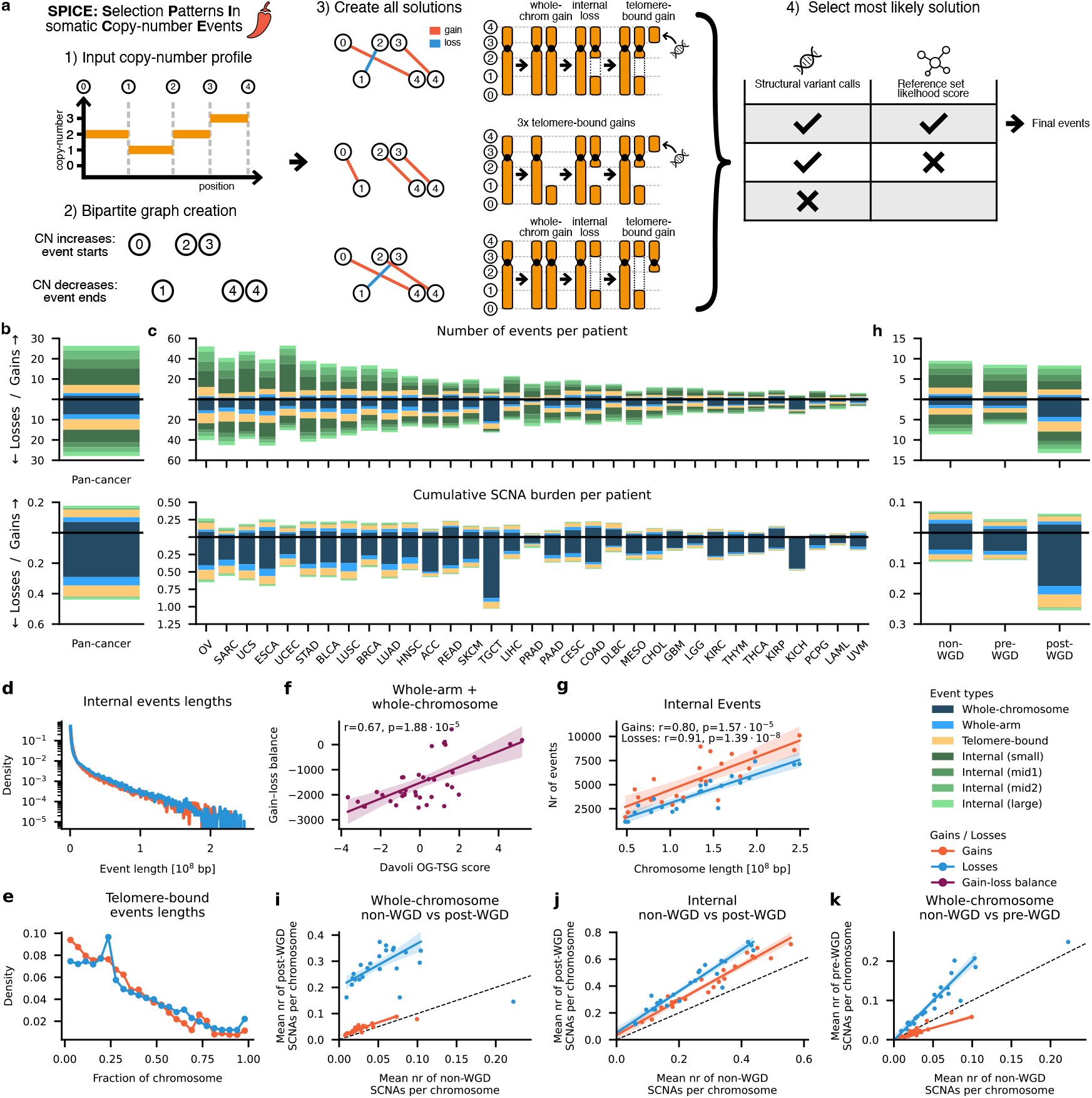
Event-level inference and global landscape of somatic copy-number alterations. **a**, Overview of the SPICE event inference framework. **b**, Composition of all somatic copy-number alteration events across event types in the pan-cancer cohort. The upper panel shows the cumulative SCNA burden per patient computed as the sum of inferred event lengths divided by the haploid genome length. **c**, Composition of SCNA event types across 33 cancer types. **d-e**, Length distribution of internal (d) and telomere-bound events (e). **f**, Correlation between the gain–loss balance (number of gains minus number of losses) per chromosome arm for whole-chromosome and whole-arm events and OG-TSG scores. **g**, Correlation between chromosome length and the number of internal events per chromosome. **h**, Composition of SCNA event types for samples grouped by whole-genome duplication (WGD) status and timing (non-, pre-, and post-WGD). **i-k**, Relative number of copy-number events per chromosome comparing non-WGD, pre-WGD and post-WGD. TCGA cancer type abbreviations are provided in the Methods.

Out of the 157,214 individual chromosomes, 54% (84,838) had a single minimum-evolution solution, while the remaining 46% (72,376) had multiple possible solutions. To resolve these ambiguous cases, we used a high-quality reference set derived from the Pan-Cancer Analysis of Whole Genomes (PCAWG) project [17, 18], which includes both allele-specific copy-number and consensus structural variant (SV) calls. For PCAWG samples, we first created candidate solutions as described above and subsequently filtered these using the available SV data, matching gains with duplications and losses with deletions to identify the most strongly supported configurations. The resulting set of unambiguous PCAWG chromosomes was then used to create the high-quality reference set of solutions. For TCGA samples, which lack SV information, and for PCAWG cases where SVs did not fully resolve the ambiguity, we selected the most plausible solution based on a similarity score to this reference set. For each ambiguous chromosome, the score was computed from event type and length, and the solution with the highest overall similarity was retained (Supplementary Figure 1; Methods).

### Patterns of somatic copy-number alterations

We detected 363,532 distinct SCNA events (median per patient = 47.0 [IQR: 26–83]; Supplementary Figure 2; Supplementary Figure 3a,b). Events were classified as whole-chromosome, whole-arm, telomere-bound, or internal (the latter subdivided into small <1 Mbp, mid1 = 1–2.5 Mbp, mid2 = 2.5–10 Mbp, and large >10 Mbp; Methods). Over 61% of events were internal (35% gains, 26% losses), and losses outnumbered gains overall—particularly for whole-chromosome alterations (Figure 1b). The median cumulative SCNA burden was 54.5% genome-equivalents (computed as the sum of inferred event lengths divided by the haploid genome length; overlapping events are counted multiple times, so values per sample can exceed 100%; Supplementary Figure 3c,d), with whole-chromosome events accounting for the majority of the cumulative SCNA burden (median of 18.0% for losses and 4.6% for gains). Thus, although small-scale events were more numerous, overall aneuploidy was dominated by large-scale alterations.

We observed substantial heterogeneity across cancer types in the number of SCNA events per patient, ranging from a median of 12.0 in Kidney Chromophobe to 93.0 in Uterine Carcinosarcoma. In contrast, the relative proportions of event types were largely consistent across cancer types, suggesting similar underlying mechanisms but differing levels of genomic tolerance (Figure 1c; Supplementary Figure 3e). Two notable outliers were kidney chromophobe (KICH) and testicular germ cell tumours (TCGT), both showing high rates of whole-chromosome losses, likely reflecting their elevated whole-genome duplication frequency (see below).

We compared our event types to published copy-number signatures [14, 15] by correlating their per-patient frequencies (Methods). We found no direct one-to-one correspondence between our event types and existing copy-number signatures—neither between our framework and those of Steele or Drews, nor between the Steele and Drews signatures themselves, indicating that each framework captures distinct mutational processes (Supplementary Figure 4). However, several meaningful associations emerged: whole-chromosome gains and losses correlated with missegregation-related Drews signatures (CX1: ***r*** = 0.19 and 0.16, respectively; CX6: ***r*** = 0.15/0.36) and WGD-associated Steele signatures (CN2: ***r*** = 0.17/0.39; CN14: n.s./***r*** = 0.19), while internal gains showed the strongest link to the homologous recombination deficiency-related signature CX5 (***r*** = 0.46).

For every gene and event type, we compared per-patient SCNA event occurrences between samples where the gene was mutated and non-mutated using a Mann-Whitney U test (corrected for multiple testing), identifying 131 genes whose mutation is associated with increased SCNA burden (Methods; Supplementary Figure 5). As expected, mutations in *TP53* and *PPP2R1A* broadly increased all copy-number event types, consistent with their roles in DNA damage repair and WGD [33, 34]. *BRCA1/2* mutations specifically upregulated telomere-bound and large internal events, likely due to homologous recombination deficiency [8]. In contrast, mutations in *CDKN2A* (cell-cycle suppression [35]) and *NOTCH1* (DNA damage response suppression [36]) were linked primarily to whole-arm and whole-chromosome alterations.

Internal event lengths ranged from 10^3^ to 10^8^ bp, with median sizes of 1.3 Mbp for gains and 0.96 Mbp for losses (Figure 1d). Event frequency decreased with length, consistent with previous reports [5, 6], and followed a Pareto distribution (Supplementary Figure 6a-d), indicating a heavy-tailed process with many short and few large events (Methods). The Pareto fit showed excellent agreement for gains (***R***^2^ = 0.99) and a good fit for losses (***R***^2^ = 0.81).

Losses tended to be shorter than gains, suggesting that partial deletions are often sufficient to inactivate genes, whereas gains require larger amplifications. Telomere-bound event lengths (expressed as a fraction of chromosome length) followed an approximately linear histogram: the shortest-length bin (0% normalized length) had frequency 1.0, and frequency declined steadily to 0.0 by 100% normalized length, consistent with uniform breakpoint generation tempered by negative selection [22] (Figure 1e).

Median event lengths were consistent across chromosomes (internal: 1.17 ± 0.29 Mbp; telomere-bound: 0.24 ± 0.05 Mbp) and cancer types (internal: 1.13 ± 0.50 Mbp; telomere-bound: 0.23 ± 0.06 Mbp) (Supplementary Figure 6e-h), suggesting that the mutational processes generating these events are broadly similar across contexts.

To assess chromosome-level selection, we compared the gain–loss balance (calculated as the number of gains minus the number of losses) of copy-number events with the oncogene–tumour suppressor gene (OG–TSG) score from Davoli *et al*. [37] (Methods). As previously reported [21], whole-arm and whole-chromosome events showed strong positive correlations with OG–TSG scores (whole-arm: ***r*** = 0.36, *p* = 0.05; whole-chromosome: ***r*** = 0.59, *p* = 2 × 10^−4^; combined: ***r*** = 0.67, *p* = 1 × 10^−5^; Figure 1f, Supplementary Figure 7a), indicating strong selection for large-scale gains in oncogene-rich regions and losses in TSG-rich regions. In contrast, smaller internal and telomere-bound events showed no such correlation, and internal event counts scaled with chromosome length (***r*** = 0.81−0.91, *p* < 10^−5^), suggesting uniform formation across the genome (Figure 1g; Supplementary Figure 7b). This behaviour was consistent across all length scales, WGD status, and for most cancer types (Supplementary Figure 7c-e).

To further test whether internal events follow a uniform genome-wide breakpoint process, we compared observed and expected event counts within common fragile sites—regions prone to breakage under replication stress in vitro [38, 39]. Expected counts were derived by randomizing fragile site positions across the genome (Methods). Only 17 of 108 sites (15.7%) showed significantly elevated event rates (*>* 1.5× expected; Supplementary Figure 7f), which can be explained by heightened selection as 16 of the 17 sites contained a known COSMIC oncogene or tumour suppressor [40]. These results suggest that although fragile sites may elevate local breakpoint formation, this alone is generally insufficient to generate a detectable enrichment in the cohort; instead, prominent signals typically require additional selection acting on nearby oncogenes or tumour suppressors, consistent with a largely uniform breakpoint process for detectable events.

### The impact of whole-genome duplication on copynumber formation

Whole-genome duplication (WGD) doubles the entire chromosome complement, amplifying chromosomal instability, buffering deleterious mutations, and accelerating clonal evolution [12, 30, 41, 42]. By modelling WGD explicitly during event inference, we partitioned events into non-WGD, pre-WGD, and post-WGD categories.

Consistent with prior reports [12], post-WGD profiles showed a pronounced shift toward losses, especially whole-chromosome losses, relative to non-WGD and pre-WGD (Figure 1h). Pre-WGD gains, by contrast, decreased modestly, likely suggesting that WGD is less advantageous in samples with already increased ploidy. Event counts remained strongly correlated across WGD states (mean ***r*** = 0.86; Figure 1i,j,k; Supplementary Figure 8), indicating that the same mutational and selective forces operate before and after WGD, as recently reported for chromosome arm-level events [42] (Methods).

The most pronounced changes occurred in post-WGD losses, which increased substantially compared to both non-WGD and pre-WGD samples (Figure 1i; Supplementary Figure 8). While most event types scaled linearly across WGD categories, whole-chromosome losses—and to a lesser extent whole-arm losses—showed a uniform, chromosome-wide increase, likely reflecting genome reduction from 4N toward the observed median ploidy of 3.3 [30]. Post-WGD samples also exhibited elevated gains across nearly all event types, consistent with heightened chromosomal instability and ongoing copy-number evolution following WGD (Figure 1j; Supplementary Figure 8) [43]. In contrast, whole-arm, telomere-bound, and internal events showed no major differences between non-WGD and pre-WGD samples, consistent with both groups lacking a completed WGD event (Supplementary Figure 8). A notable exception is the increase in whole-chromosome losses in pre-WGD relative to non-WGD samples, suggesting that such losses may precede WGD (possibly through the generation of widespread loss of heterozygosity (LOH) regions, Supplementary Figure 8) and might contribute to triggering WGD. Additionally, pre-WGD samples showed fewer whole-chromosome gains than non-WGD samples, potentially reflecting selection against high pre-duplication ploidy, or selection against WGD in samples with more whole-chromosome gains (Figure 1k).

### A simple selection model can explain the observed copy-number landscape

We developed a generative model of somatic copy-number selection that operates directly on individually inferred internal SCNA events to identify loci under selection (Figure 2a; Supplementary Figure 9; see Methods for a detailed description). Unlike previous approaches that detect peaks in aggregated copy-number signals, our model operates directly on inferred events and treats genome-wide breakpoint formation as a neutral reference against which locus-specific alterations are detected. We focused on internal SCNAs, for which the assumption of uniform breakpoint formation is most appropriate. For each chromosome, we summarized the data as eight event tracks defined by SCNA polarity (gain or loss) and the four event-length classes (small, mid1, mid2, large; Supplementary Figure 9a). Candidate loci are defined by a genomic position and a selection strength for each track. In the generative process, the observed events are randomly reassigned along the chromosome under uniform breakpoint formation, and events overlapping a selected locus are preferentially retained there in proportion to the track-specific selection strength of the locus. This produces expected event-count profiles that combine a flat baseline with triangular enrichments centred on selected loci, with triangle width determined by event length and height by selection strength (Supplementary Figure 9b). For efficiency, explicit event-level simulations are replaced by an equivalent convolution-based implementation (Supplementary Figure 9c).

**Figure 2.**
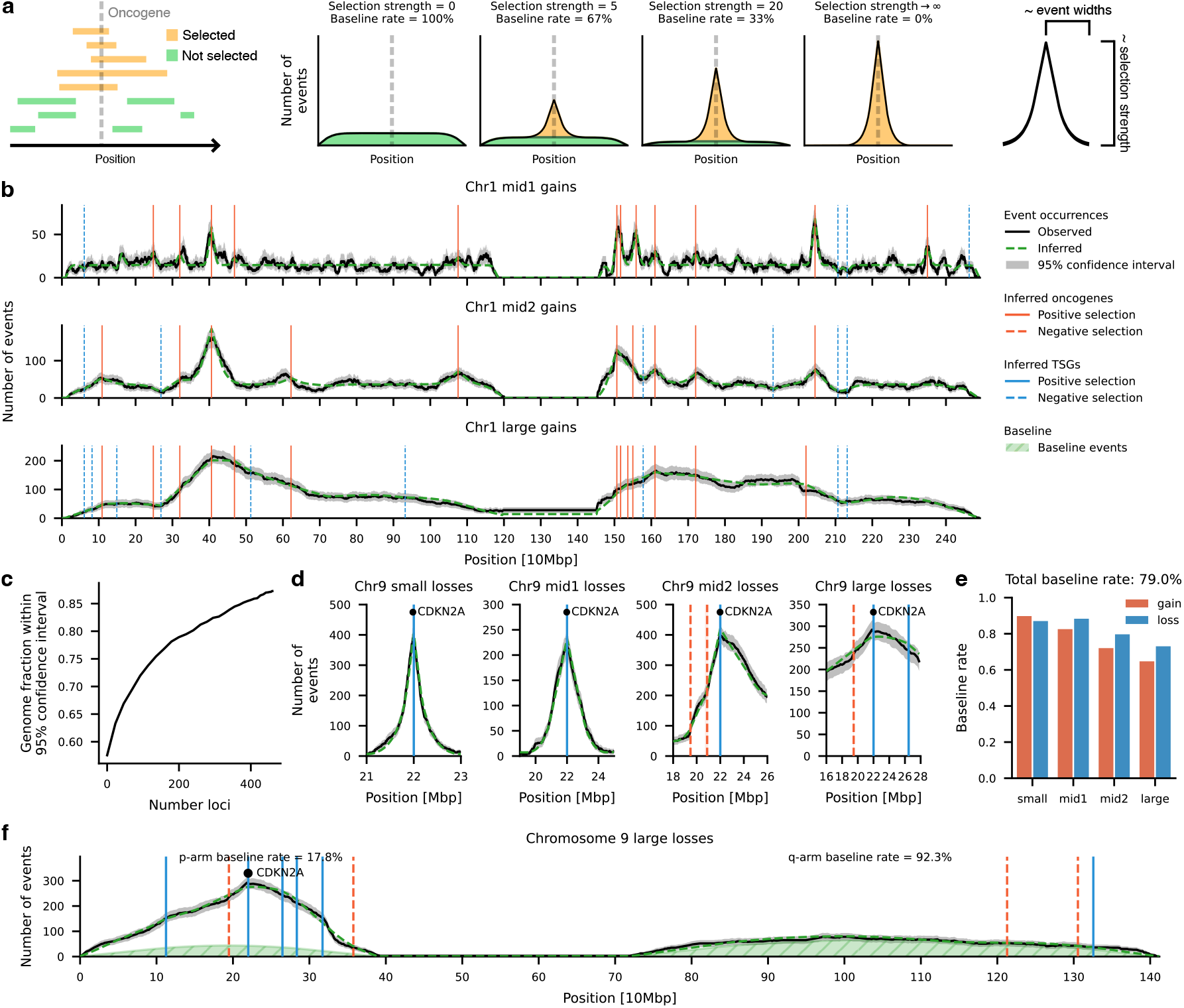
Generative modelling of focal selection in somatic copy-number events. **a,** Overview of the SPICE generative selection model. **b**, Example model fits for internal gains on chromosome 1 across three length classes (mid1, mid2, large). **c**, Fraction of the genome whose observed signal lies within the model’s 95% confidence interval as the number of selected loci increases. Loci are ordered by the order they were added in the MCMC approach. **d**, Model fits for losses overlapping the tumour-suppressor locus *CDKN2A* on chromosome 9 across four internal event length classes. **e**, Baseline rate per event class, showing the proportion of internal gains and losses not assigned to loci under selection. **f**, Example of inferred loci and baseline rates for the parm q-arm of chromosome 9. Note that by design, there are no events in the centromere region between the two arms.

Loci under selection are not specified a priori but are inferred by fitting the generative model to the observed event tracks. We used an iterative Markov Chain Monte Carlo procedure that proposes, repositions, and rescales loci so as to minimise the mean squared error between observed and model-generated event counts across the chromosome across all eight tracks (Methods; Supplementary Figure 9d). Each locus is classified as oncogene-like or tumour-suppressor-like, such that oncogene-like loci increase selection for gains and decrease it for losses, whereas tumour-suppressor-like loci have the opposite effect. A final post-processing step merges nearby loci representing the same signal and removes loci with weak support, yielding a parsimonious set of selected loci whose combined effect explains the observed focal copynumber landscape.

Applied to the 5,966 TCGA samples (Supplementary Figure 10), the model identified 460 loci under selection (285 oncogene-like, 175 tumour-suppressor-like; Figure 2b; Supplementary Figure 11a). Locus significance was assessed by permutation against the neutral shuffling model, with Benjamini-Hochberg correction and a cutoff of *p* < 0.05 (Methods). Longer chromosomes contained more loci (***r*** = 0.75, *p* = 3.5 × 10^−5^ for gains and ***r*** = 0.71, *p* = 1.5 × 10^−4^ for losses; Supplementary Figure 11b,c). Despite the simplicity of our model, simulated and observed event counts agreed closely, with 89.3% of the genome within the 95% confidence interval of the observed signal (Methods; Figure 2c; Supplementary Figure 11d).

The characteristic triangular enrichment patterns predicted around selected loci were consistently observed across the TCGA data (Supplementary Figure 12). These are best exemplified by the well-characterized tumour suppressor *CDKN2A*, where the model accurately reproduces the observed signal for 1,025 unique losses (Figure 2d). In the mid2 length scale, *CDKN2A* events showed a slight right-shift in observed counts due to negative selection from nearby OG-like loci (*ACER2, MLLT3*), which the model captures accurately, demonstrating the ability to model both positive and negative selection.

### The majority of internal copy-number events are not subject to focal selection

Using our model assumptions, each observed event can probabilistically be assigned to the loci under selection or be classified as a baseline event. In practice, events not overlapping any locus of selection were classified as baseline events, while overlapping events were assigned a baseline probability inversely related to the cumulative selection strength of overlapping loci (Figure 2a; Methods). The overall baseline rate of a given chromosome is then obtained by aggregating the baseline probabilities across all its events.

The landscape of SCNA events not under selection had previously not been comprehensively mapped and our approach established a total baseline rate of 79.0%, indicating that the vast majority of internal SCNAs are not subject to focal selection (Figure 2e; Supplementary Figure 13a). The behaviour of baseline events is best exemplified by losses on chromosome 9, where the p-arm contains multiple strong TSGs, most notably *CDKN2A*, and consequently harbours only few baseline losses (17.8%). By contrast, the p-arm contains only a single weak TSG and subsequently is dominated by the presence of baseline losses (92.3%) and is therefore perfectly modelled through the uniform baseline distribution (Figure 2f). Overall baseline rates varied by chromosome, ranging from 54.0% for losses on chromosome 9 to 95.6% for gains on chromosome 21, largely reflecting the presence of strong OG and TSG loci (Supplementary Figure 13b-d). The baseline rate was highest for small events (89.8% and 87.0% for gains and losses, respectively) and lowest for large events (64.7% and 73.0% for gains and losses, respectively), consistent with the expectation that larger events are more likely to overlap multiple genes and be subject to selection. We found baseline rates to be stable across WGD status and cancer types (Supplementary Figure 13e,f). Finally, to confirm that baseline rates were not artefacts of our event inference, we examined breakpoint distributions in the raw copy-number profiles. The uniformity of break-point spacing per chromosome negatively correlated with inferred baseline rate (***r*** = −0.55, *p* = 0.0071), indicating that chromosomes with higher baseline rates indeed showed more uniformly distributed breakpoints (Methods; Supplementary Figure 13g,h).

### SPICE identifies hundreds of novel loci under SCNA selection

We identified 460 loci under selection—substantially more than reported by previous methods at the same p-value threshold (*p* < 0.05; 110 by GISTIC [6] and 134 by BISCUT [22]). We recovered most previously reported loci (72/110 GISTIC, 69/134 BISCUT; Figure 3a; Supplementary Figure 12a; Supplementary Figure 14a; Supplementary Figure 15), with overlap significance estimated by a permutation test that randomly repositions loci across the genome (*p* < 0.0001 for both; Methods). Of the GISTIC loci not detected by SPICE, 10 were within 1 Mbp of one of our loci, likely reflecting overly strict locus boundaries, and another 18 were within 2 Mbp of centromeres or telomeres, suggesting boundary-related limitations (Supplementary Figure 14b-d). For BISCUT, discrepancies likely arise from methodological differences, as BISCUT was trained on telomere-bound rather than internal events. Around the missed BISCUT loci, our model explained 87.6% of the genome within 95% confidence intervals without requiring loci under selection (Supplementary Figure 14e,f).

**Figure 3.**
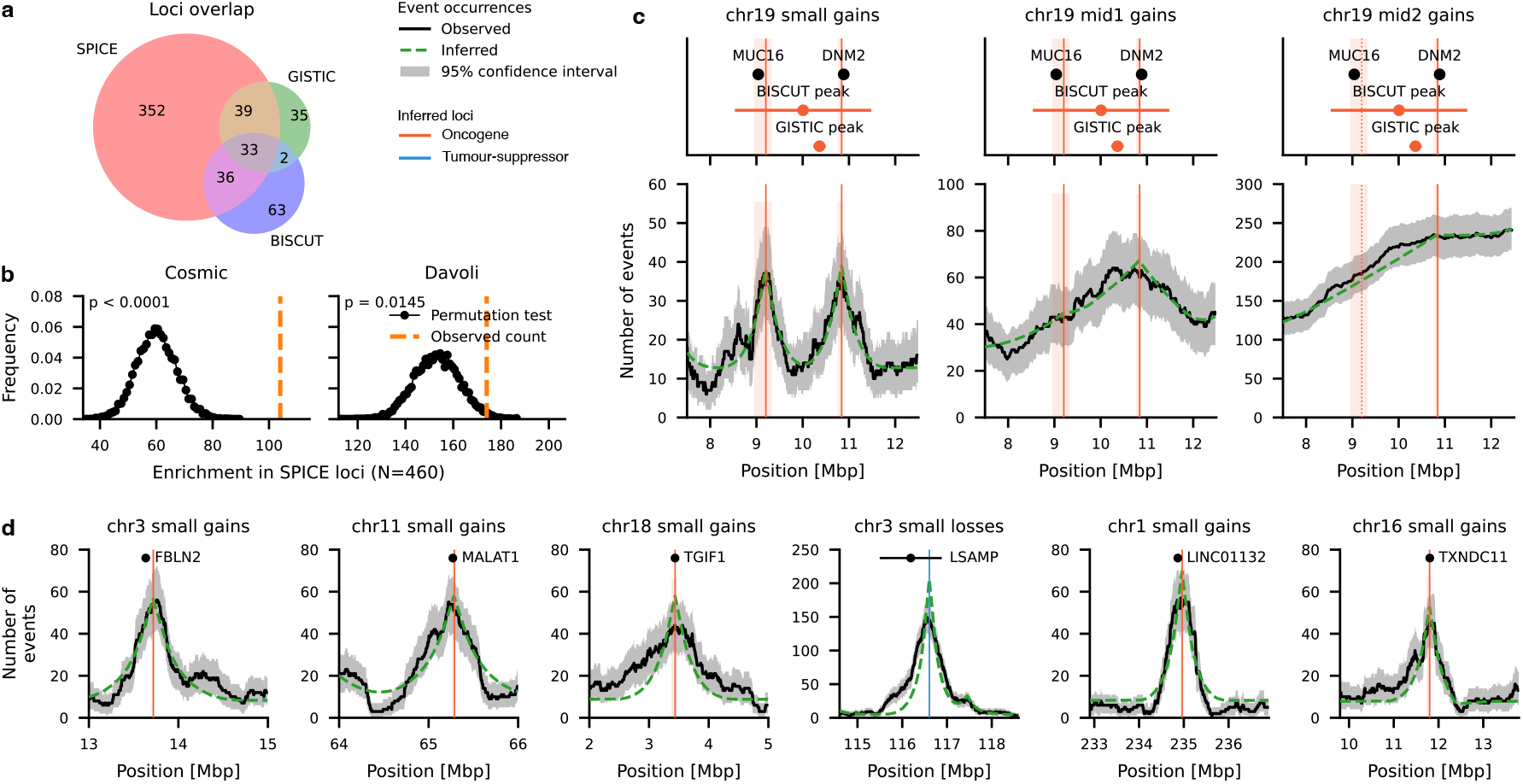
SPICE identifies loci under somatic copy-number selection. **a**, Overlap of loci identified by SPICE, GISTIC, and BISCUT showing the number of shared and unique loci among methods. **b**, Enrichment of the 460 SPICE loci for known cancer genes using COSMIC annotations (left) and mutation-based driver scores from Davoli *et al*. (right) using a permutation test. **c**, Comparison of SPICE with GISTIC and BISCUT for gains on chromosome 19 at three event length scales (small, mid1, mid2). Shown are the positions of loci reported by each method and known oncogenes (*MUC16, DNM2*), along with observed event counts, model predictions, and confidence intervals, illustrating how analysis across multiple length scales resolves overlapping signals that were merged or misplaced by previous methods. **d**, Examples of novel loci identified by SPICE, not detected by previous SCNA methods, including *FBLN2* (chr3), *MALAT1* (chr11), *TGIF1* (chr18), *LSAMP* (chr3), *LINC01132* (chr1), and *TXNDC11* (chr16). Each panel shows observed and modelled event counts across the locus.

To further validate our inferred loci, we tested for enrichment of known cancer genes. 104 SPICE loci contained at least one COSMIC gene [40]—nearly twice the expected number from the permutation test (observed-to-expected ratio = 1.75; *p* < 0.0001; Figure 3b; Methods). Using mutation-based driver scores from Davoli *et al*. [37], we found that 174 loci contained a significant oncogene or tumour suppressor (observed-to-expected ratio = 1.14; *p* = 0.015; Figure 3b). Enrichment levels were comparable to those achieved by GISTIC and BISCUT (ratio observed/expected for GISTIC: 1.90 and 1.25, BISCUT: 1.51 and 1.18 for enrichment in COSMIC and Davoli genes, respectively; Supplementary Figure 14g), supporting the biological relevance of our discoveries.

In addition to recovering most previously reported loci, we identified 352 novel regions under selection not detected by either GISTIC or BISCUT (Supplementary Figure 12b). SPICE’s enhanced sensitivity stems partly from its use of multiple length scales, which helps resolve overlapping signals that obscure distinct loci (Supplementary Figure 16). On chromosome 19 near 10 Mbp, both GISTIC and BISCUT report a single locus, but small-scale analysis reveals two separate loci under selection that merge at larger scales (Figure 3c; Supplementary Figure 16a). As a result, BISCUT merges the two loci into one, while GISTIC places its locus incorrectly between them. The first locus contains the COSMIC oncogene *MUC16* and the second locus the COSMIC oncogene *DNM2*. A similar pattern occurs on chromosome 1 near the centromere (150–162 Mbp), where small-scale analysis resolves six loci from two GISTIC loci (Supplementary Figure 16b), five of which contain known oncogenes, confirming that SPICE identifies additional true driver regions missed at coarser resolutions.

Among the novel loci, *FBLN2* (Tier 2 COSMIC oncogene) showed recurrent gains and is overexpressed in urothelial and lung adenocarcinoma [44, 45] (Figure 3d; Supplementary Figure 17). Another locus contains *MALAT1*, a long non-coding RNA with known oncogenic activity across multiple cancer types [46]. We also find a locus at *TGIF1* on chromosome 18, previously classified only as a tumour suppressor [37], but recurrently gained in our data, consistent with reports of oncogenic roles in glioma, breast, and lung cancer [47, 48].

Finally, 196 loci were not reported by any previous method and did not contain any known TSG or OG from COSMIC or Davoli *et al*. These include a tumour-suppressive locus at *LSAMP* on chromosome 3, which is thought to facilitate epithelial-mesenchymal transition [49] (EMT; Figure 3d; Supplementary Figure 17), an oncogenic locus at *LINC01132* overexpressed in hepatocellular carcinoma [50], and an OG locus on chromosome 16 containing *TXNDC11* and *BCAR4*, both implicated in multiple cancers and therapy resistance [51–53].

### Cancer type–specific patterns of selected loci

Using the event-to-locus assignments, we counted the number of events linked to each locus in every sample. For every cancer type, we calculated both the fraction of samples harboring at least one event overlapping a given locus and the mean number of events per sample. These aggregated values were used to compare selection patterns across tumour types (Figure 4a, Supplementary Figure 18).

**Figure 4.**
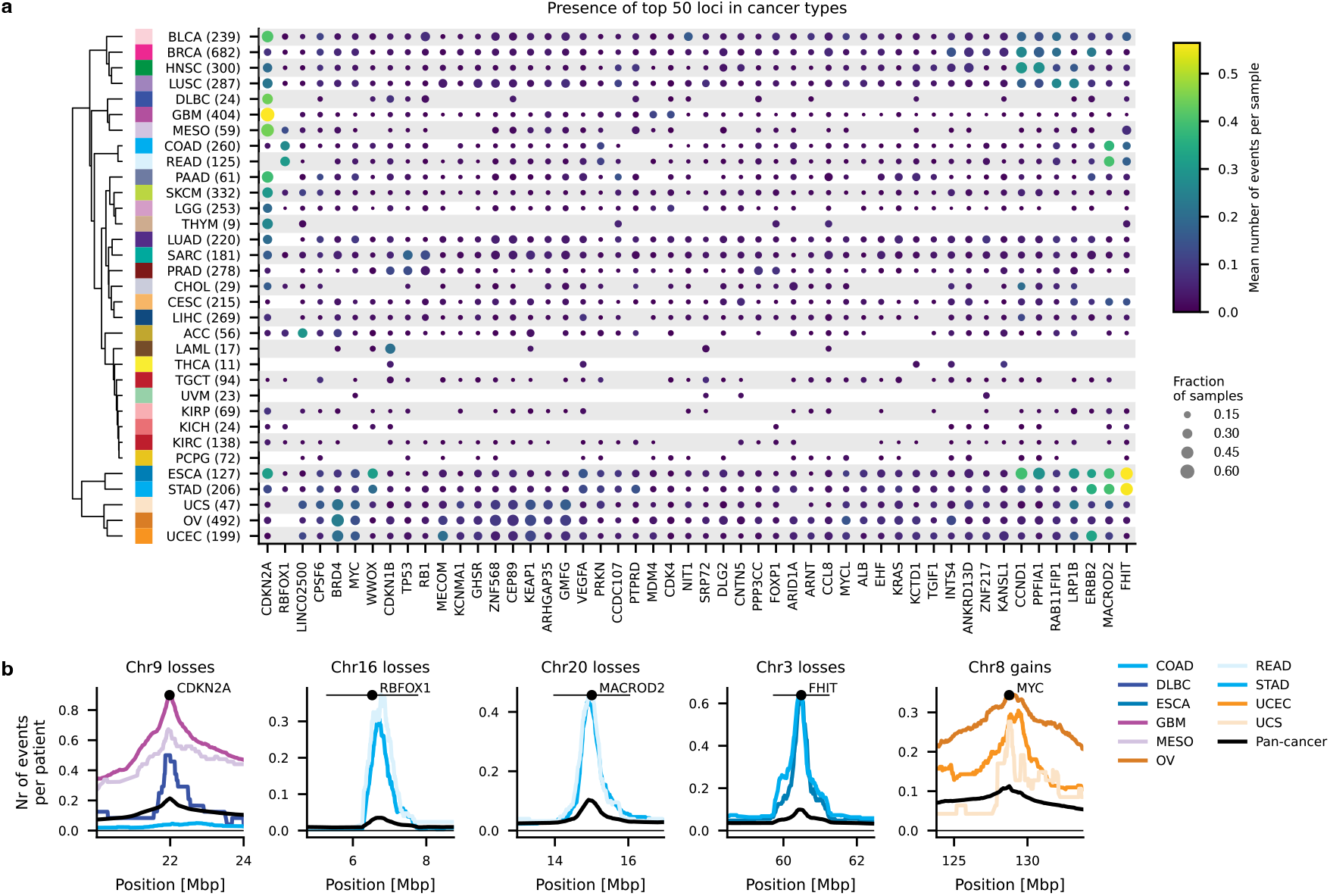
Cancer type–specific patterns of loci under somatic copy-number selection. **a**, Presence of the top 50 loci under selection across 33 cancer types. Loci are ranked by the total number of events assigned to them and clustered by their occurrence profiles. Cancer types are hierarchically clustered based on selection patterns across all loci. Dot size indicates the fraction of samples within a cancer type containing at least one event overlapping a given locus, and color represents the mean number of events per sample. **b**, Examples of cancer type–specific selection profiles showing the number of events per patient across loci. Shown are losses at *CDKN2A* (chr9), *RBFOX1* (chr16), *MACROD2* (chr20), and *FHIT* (chr3), and gains at *MYC* (chr8), illustrating differences in locus-level selection strength and tumour-type specificity. TCGA cancer type abbreviations are provided in the Methods.

*CDKN2A* exhibited frequent copy-number alterations across several cancer types, including glioblastoma, mesothelioma, and diffuse large B cell lymphoma, while being largely absent in others such as colorectal cancers, illustrating the tumour-type specificity of locus-level selection patterns (Figure 4b). Hierarchical clustering of cancer types based on these locus-level occurrence profiles revealed distinct groupings. Several cancer-type clusters showed enrichment for shared loci. For example, colorectal cancers (COAD, READ) grouped together with strong selection at *RBFOX1*, which has recently been shown to undergo frequent focal deletions and reduced expression in colorectal cancer [54], and at *MACROD2*, a mono-ADP-ribosylhydrolase frequently deleted in gastrointestinal and other malignancies [55]. Upper gastrointestinal cancers (ESCA, STAD) showed focal selection for example at *FHIT*, a candidate tumour suppressor located within the FRA3B fragile site [56]. Estrogen-dependent gynecologic tumours (OV, UCEC, UCS) clustered via selection at multiple loci such as *MYC*, which is involved in the tumourigenesis of a wide range of cancer types [57].

## Discussion

Here, we introduce SPICE, a framework for inferring discrete copy-number events and identifying loci under selection from allele-specific copy-number profiles. SPICE overcomes the ambiguity of copy number profiles and reconstructs the discrete events changing the genome. SPICE employs a minimum-evolution procedure for event reconstruction that accounts for whole-genome duplication (WGD). Using this approach, we inferred 363,532 somatic copy-number alteration (SCNA) events across 5,966 TCGA samples covering 33 cancer types. The analysis revealed distinct cancer-type-specific and WGD-dependent patterns in event distributions. We further showed that internal events arise largely from a uniform breakpoint process shaped by focal selection.

To model SCNA selection, we developed a generative framework that explains the vast majority of the observed copy-number landscape characterised by triangular shapes around loci under selection. Working directly from inferred events rather than aggregated copy-number profiles, SPICE identified 460 loci under selection—recovering most previously inferred SCNA tumour suppressors and oncogenes from GISTIC and BISCUT and revealing 352 novel loci. Although loci under selection were inferred in a pan-cancer cohort, we found strong differences in locus strength and occurrence across cancer types. Notably, the model does more than find peaks: it explains most of the genome with a simple, dosage-based selection mechanism in which loci exert approximately additive fitness effects on gains and losses. In this view, more complex local patterns emerge naturally from the superposition of nearby oncogene-like and tumour-suppressor-like loci, and resolve clearly only in large cohorts. This suggests that much of copy number selection can be captured by a relatively small set of locus-level dosage effects.

Most internal events (79.0%) could be explained by a baseline, approximately uniform breakpoint-formation process across the genome, rather than by focal selection. Consistent with this, only a minority of common fragile sites showed elevated internal event rates, largely attributable to nearby driver loci; however, we cannot exclude that this estimate is biased by cancer-type-specific fragile-site activity. As these baseline events are not under positive selection, they represent the SCNA equivalent of passenger mutations in point mutation studies [58]. The high baseline rate across the whole genome contrasts with previous results based only on telomere bound events which suggested that cancer aneuploidies are primarily shaped by fitness effects [22]. This discrepancy may reflect distinct evolutionary dynamics between internal and telomere-bound events, as the latter could be more disruptive and thus contain fewer neutral events. Alternatively, selection-neutral telomere-bound events may simply be harder to detect. A further confounder is that the baseline rate will be inflated if too few loci are inferred, or if true loci are removed because they do not pass the *p* = 0.05 threshold; the true baseline rate is therefore likely somewhat lower. However, because the generative model still fits the observed landscape closely (89.3% of the genome within the 95% confidence interval), this inflation is unlikely to be large, and the conclusion of a high baseline rate remains.

By modelling across multiple length scales, SPICE improves the localization of driver loci and reveals focal regions obscured at coarser resolutions. It advances previous methods by using a generative model grounded in simple evolutionary principles to detect tumour-suppressor and oncogene loci—unlike GISTIC, which identifies peaks in alteration frequencies [6, 25], or BISCUT [22], which relies on breakpoint distributions. The model’s close fit to observed data supports a view of SCNA selection as driven by uniform breakpoint formation, focal selection, and a high baseline rate.

At the same time, SPICE has limitations. It assumes approximately uniform breakpoint formation, which holds for most internal events but not for telomere- or centromere-anchored events, so these were excluded from the current study. Sensitivity near chromosome ends and centromeres is reduced, yet strong loci such as *FGFR3* on chromosome 4 (1.78 Mbp from the telomere) are still detected. Inferring loci under selection also requires large sample sizes, as triangular loci shapes emerge only with sufficient data. Even with thousands of tumours, some loci remain weak, making de novo discovery in smaller datasets infeasible. However, as copy-number studies expand, the number of high-quality samples will continue to increase. Finally, some cancers follow non-parsimonious evolutionary routes, though these typically involve only a single additional event [42].

In future work, event reconstruction could be improved by replacing the linear genome with a graph-based representation, such as Junction Balance Analysis (JaBbA) graphs, which encode rearranged chromosomes [11]. Additionally, recent studies have used point mutations to time SCNAs [42], which could help refine event ordering. We also aim to experimentally validate the top novel loci, apply the framework to singlecell copy-number data, and integrate structural-variant, expression, and functional screening data to refine locus selection models.

In summary, SPICE provides a comprehensive framework for dissecting the evolutionary forces acting on somatic copynumber alterations, revealing how selection shapes oncogene activation and tumour-suppressor loss across thousands of cancer genomes.

## Methods

### Data collection and pre-processing

We downloaded allele-specific copy-number profiles as well as whole-genome duplication (WGD) status for 10,674 samples from The Cancer Genome Atlas (TCGA) from the AS-CAT v3 repository [26]. Preprocessing was done using CNSistent [27] and our custom pipeline. It included imputing missing segments from the neighbouring segments, capping copynumber values at 8 and removal of small segments (<1kbp). Phasing was carried out using MEDICC2, which assigns major and minor alleles to pseudo-alleles A and B to minimize the total number of events from a diploid karyotype [29]. We filtered the dataset to 6,335 high-quality samples using the whitelist from Drews *et al*., which excludes noisy profiles, low-quality arrays, and samples lacking detectable chromosomal instability. In studies with multiple samples per tumour, the whitelist selected a single representative [15]. Samples with multiple WGDs were furthermore excluded, resulting in a final cohort of 5,966 samples.

Copy number profiles of samples were split into individual allele-specific chromosomes (hereafter simply referred to as chromosomes) and removed chromosomes without any SC-NAs. Each chromosome was processed independently. We excluded highly re-arranged chromosomes with more than 40 inferred events (based on the MEDICC2 event distance [29]), as these likely reflect complex, non-parsimonious rearrangement processes for which our inference assumptions may not hold [10, 11], and discarded chromosomes that failed processing due to excessive loss of heterozygosity (see below).

TCGA data was used for the primary analysis due to its large sample size and broad cancer type coverage. In addition, we used a secondary dataset from the Pan-Cancer Analysis of Whole Genomes (PCAWG) [17] as a reference dataset (see below), selected for its high-quality copy-number profiles and available structural variant calls. We used the PCAWG allele-specific consensus copy-number calls and consensus WGD assignments [18] and applied the same preprocessing steps as above, yielding 2,543 samples and 57,188 non-neutral chromosomes.

## TCGA cancer type abbreviations

### Inference of events

For a proof of SPICE’s event inference, see the Appendix.

Inference of copy-number events using bipartite graphs Copy-number events were inferred using our previously established minimum evolution framework [28, 29]. First, each non-neutral chromosome’s copy-number profile was converted into a list of integers and padded with the neutral value for allele-specific chromosomes at both ends—1 if no WGD was detected and 2 if WGD was present. We defined a copy-number gain as an increase over a contiguous region, introducing two breakpoints: one where the copy-number rises and one where it falls. A loss is analogously defined, with break-points where it drops and then rises.

Using this, we encoded the breakpoints of each profile as a bipartite graph: one set of nodes for rising breakpoints, one for falling breakpoints (Supplementary Figure 1b). If a change of magnitude k occurs at breakpoint b, we add k nodes labelled b to the appropriate set. As each copy-number event corresponds to two breakpoints (one dropping and one rising), events are represented as edges connecting a pair of nodes (one from each set). If the rising breakpoint precedes the falling one, it represents a gain; otherwise, a loss. Padding ensures events that are telomere-bound or span entire chromosomes are captured correctly. Multiple valid pairings represent different possible event histories, all with the same number of events.

**Table 1.** TCGA study abbreviations used in this work.

| Study abbreviation | Study name |
| --- | --- |
| LAML | Acute Myeloid Leukemia |
| ACC | Adrenocortical carcinoma |
| BLCA | Bladder Urothelial Carcinoma |
| LGG | Brain Lower Grade Glioma |
| BRCA | Breast invasive carcinoma |
| CESC | Cervical squamous cell carcinoma and endocervical adenocarcinoma |
| CHOL | Cholangiocarcinoma |
| COAD | Colon adenocarcinoma |
| ESCA | Esophageal carcinoma |
| GBM | Glioblastoma multiforme |
| HNSC | Head and Neck squamous cell carcinoma |
| KICH | Kidney Chromophobe |
| KIRC | Kidney renal clear cell carcinoma |
| KIRP | Kidney renal papillary cell carcinoma |
| LIHC | Liver hepatocellular carcinoma |
| LUAD | Lung adenocarcinoma |
| LUSC | Lung squamous cell carcinoma |
| DLBC | Lymphoid Neoplasm Diffuse Large B-cell Lymphoma |
| MESO | Mesothelioma |
| OV | Ovarian serous cystadenocarcinoma |
| PAAD | Pancreatic adenocarcinoma |
| PCPG | Pheochromocytoma and Paraganglioma |
| PRAD | Prostate adenocarcinoma |
| READ | Rectum adenocarcinoma |
| SARC | Sarcoma |
| SKCM | Skin Cutaneous Melanoma |
| STAD | Stomach adenocarcinoma |
| TGCT | Testicular Germ Cell Tumors |
| THYM | Thymoma |
| THCA | Thyroid carcinoma |
| UCS | Uterine Carcinosarcoma |
| UCEC | Uterine Corpus Endometrial Carcinoma |
| UVM | Uveal Melanoma |

Out of the 157,214 non-neutral chromosomes identified from TCGA, 84,838 chromosomes (54.0%), had only a single solution. For 63,273 chromosomes (40.2%) with fewer than 10 events, we enumerated all possible solutions. For the remaining 9,103 chromosomes (5.8%) with ≥ 10 events, we used a Markov Chain Monte Carlo (MCMC) approach to explore the solution space efficiently (see below).

#### Loss of heterozygosity segments

Loss of heterozygosity (LOH) occurs when one allele of a genomic segment is completely lost, resulting in an allele-specific copy number of zero. In regions without LOH, the order of gains and losses is irrelevant. In contrast, for LOH segments the event order matters, since a fully lost segment cannot be regained. The bipartite graph representation can produce invalid solutions in which a LOH segment is never fully lost or losses occur too late, leaving segments with nonzero copy number when they should be zero (Supplementary Figure 1c).

LOH segments can also introduce spurious events: if a segment is lost, its neighbors become adjacent, and subsequent gains that overlap both neighbouring segments may appear as if the LOH segment was lost twice. To prevent such artefacts, LOH segments were adjusted by setting their copy number to min(*C*_*i*−1_, *C*_*i*+1_) − 1, where *C*_*i*−1_ and *C*_*i*+1_ are the copy-number values of adjacent segments. This correction avoids spurious events inconsistent with minimum-evolution.

To ensure that all, and only, LOH segments are fully lost, we require at least one possible ordering of events in which every LOH segment reaches a copy number of zero at some point, while all non-LOH segments always remain above zero. The order of events refers to the sequence in which the events are applied to the diploid copy-number profile.

We formulated this as a constraint programming problem and solved it using Google OR-Tools’ CP-SAT solver [59]. Each bipartite-graph solution was represented as an ***N*** × ***M*** matrix, where ***N*** is the number of events in the minimum-evolution solution and ***M*** is the number of genomic segments. Matrix entries encode event overlap: gains as +1, losses as −1, and 0 otherwise. The solver searches for an event order (i.e., permutation of matrix rows) such that the cumulative sum across events—representing successive copynumber states—satisfies LOH constraints. Specifically, for each LOH segment, there must be a row where the cumulative sum of values in the column representing the segment is −1, while non-LOH segments must always remain *>* 0. Solutions with at least one valid ordering were retained; all others were discarded. The original, unadjusted profile can then be reconstructed simply by discarding all gains that would happen after −1 was reached.

#### Whole-genome duplications (WGD)

During event inference, special attention must be given to whole-genome duplications (WGD), drastic events that double the entire genomic content. We inferred the WGD status of each patient as described above. For WGD-positive patients, a single WGD event must be placed at an appropriate point in the evolutionary history, distinguishing pre-WGD from post-WGD events.

First, we constructed the bipartite graph as previously described (Supplementary Figure 1d). Due to genome doubling, the number of nodes typically exceeded the number of minimum-evolution events, as breakpoints created before WGD are duplicated. To account for this, we computed the expected number of events under the minimum-evolution model using MEDICC2’s [29] event distance with the –wgd-x2 flag enabled. From this expected number of events and the number of nodes, we inferred how many pre-WGD events must be present, and used this difference to guide subsequent analysis.

As a first step, we identified pre-WGD events as identical edges—pairs with the same start and end nodes—which were merged and treated as single pre-WGD events. The remaining nodes were then connected to represent the post-WGD events.This step is necessary because certain combinations of events—such as a pre-WGD gain followed by a post-WGD loss, or a pre-WGD loss followed by a post-WGD gain—can share a breakpoint that becomes invisible in the final copy-number profile, as the opposing events cancel each other out. By including these duplicate node pairs, we ensured that such hidden configurations were represented. The solution search was then re-run on the augmented graph, with the increased node count balanced by the corresponding number of inferred pre-WGD events.

#### Selection of the most plausible event solutions

For chromosomes with fewer than ten inferred events, all valid event solutions were enumerated, and the most plausible one was selected using a combination of structural variant (SV) support and similarity scoring against a high-quality reference set derived from PCAWG. When SV data were available (PCAWG samples), we first filtered solutions to retain only those with the maximum number of SV overlaps (Supplementary Figure 1e). When no SV calls were available (TCGA samples or incomplete PCAWG data), or when multiple equally supported solutions remained, we selected the final solution using a similarity score based on reference event profiles (Supplementary Figure 1f).

For SV-supported disambiguation, we counted the number of inferred events in each solution that overlapped a structural variant of the corresponding type—gains with duplications and losses with deletions—using the PCAWG consensus SV dataset. Solutions with the highest number of matching SV overlaps were retained, while those with fewer overlaps were discarded. In cases where this filtering step left a single remaining solution, it was directly selected as the final event configuration.

If no SV data were available or the SV information did not fully resolve the ambiguity, we applied a similarity-based approach to identify the most plausible solution. The reference set used for this procedure comprised 27,888 PCAWG chromosomes with unique solutions and an additional 5,327 chromosomes with multiple candidate solutions that could be fully resolved using SV support. This combined set was used as a high-confidence reference to guide ambiguous cases toward solutions that most closely resembled realistic event profiles observed in the reference data.

For each candidate solution, a distance-based similarity score was computed as follows. (1) Each event was assigned to one of 12 categories defined by whole-genome duplication (WGD) timing (no-WGD, pre-WGD, or post-WGD) and event type (internal, telomere-bound, whole-arm, or whole-chromosome). (2) For two events within the same category—one of length ***L***_1_ on a chromosome of length ***C***_1_, the other of length ***L***_2_ on a chromosome of length ***C***_2_—the distance was defined as 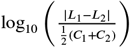. When ***L***_1_ = ***L***_*2*_ (e.g., for identical whole-arm or whole-chromosome events), it was set to −6. (3) For each event *e* in the candidate solution, we computed its distance *d*(*e, r*) to every event ***r*** in the reference set belonging to the same category and summed the 250 smallest distances to obtain a cumulative event distance. (4) The total similarity score for a solution was the sum of cumulative distances across all its events, and the solution with the lowest total score was selected.

This approach effectively models the likelihood of event lengths and types in a non-parametric manner, favoring solutions that closely match the empirical distributions of event characteristics observed in the high-confidence reference dataset.

#### MCMC-based event inference for highly rearranged chromosomes

For the 9,103 chromosomes (5.8%) containing ten or more events, exhaustive enumeration of all possible event solutions was computationally infeasible. Instead, we used a Markov Chain Monte Carlo (MCMC) approach with simulated annealing to efficiently explore the solution space.

As before, a bipartite graph was constructed and the MCMC was initialized with a random valid solution by pairing nodes, and its similarity score to the reference set was computed. At each iteration, a new proposal was generated by modifying the current solution. For non-WGD chromosomes, proposals involved swapping the end nodes of two existing edges in the bipartite graph. For WGD chromosomes, proposals included both such edge swaps and the addition or removal of pairs of nodes with identical labels. Each proposed configuration was checked for LOH validity and scored using the similarity score to the high-confidence reference set derived from PCAWG.

Acceptance of new proposals followed a simulated annealing scheme. If the current score ***S***_current_ was better than the best score ***S***_best_, the proposal was accepted. Otherwise, it was accepted with probability 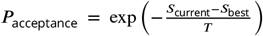. The temperature parameter ***T*** decreases over time according to ***T***_*i*_ = 10^1−5*i*/*n*^, for iteration ^*i*^ and total number of iterations ^*n*—^allowing broad exploration at high temperatures and convergence toward the optimal solution as the temperature cools.

#### Post-processing and classification of events

To improve event quality and consistency, we applied several post-processing filters. Events located within 1 Mbp of a centromere were excluded due to poor mappability in these regions. Events were classified based on their relation to chromosome ends: events containing the first or last copy-number segment were labelled as telomere-bound; events telomere-bound on both sides were defined as whole-chromosome events; and those telomere-bound on a single side and spanning 95–105% of an arm were considered whole-arm events. Events not meeting these criteria were classified as internal. To reduce artefacts near telomeres, telomere-bound events shorter than 1% of the chromosome length were discarded. Finally, duplicate events—those with identical start and end coordinates on the same chromosome of the same sample—were removed, as they likely represent amplifications or complex rearrangements.

We stratified internal events into four length classes based on their empirical size distribution per chromosome, excluding outliers smaller than 100 kbp or larger than 40 Mbp. The final classes were defined as small (<1 Mbp), mid1 (1–2.5 Mbp), mid2 (2.5–10 Mbp), and large (>10 Mbp).

#### Removal of recurrent technical artefacts

In the aggregated event signal, we observed rectangular, step-like features that we term plateaus (Supplementary Figure 19). Unlike true selection loci, plateaus arise when many samples share nearly identical breakpoints, producing sharp edges and a flat interior in the signal.

Multiple lines of evidence indicate these are technical artefacts. The strongest example is *HYDIN* on chr16, where >600 gains share the same breakpoints; this coincides with a recent duplication missing from the hg19 reference [60]. More generally, plateaus seen in TCGA data vanish in PCAWG whole-genome data, both at aggregate and per-sample levels, confirming they reflect platform and copy-number calling issues rather than biology.

We detected plateaus per chromosome and length class by identifying regions with extreme breakpoint steepness (top 1%), ≥ 1 Mbp width, ≥ 2× signal over flanks, and flat interior, defined as (max − min)/mean < 1. Candidates were manually reviewed, yielding 22 plateaus across 12 chromosomes. For each, 1 Mbp start and end regions were defined, and any event overlapping these boundaries was removed. This filtering prevented artefactual loci and improved the specificity of down-stream analyses.

### Analysis of inferred events

#### Correlation with external data

We correlated copy-number event type occurrence with copynumber signatures taken from Steele *et al*. and Drews *et al*. [14, 15]. Event type occurrences were normalized per patient (such that the occurrence per patient sums to 1). Correlations were performed using Spearman’s rank correlation, and multiple-testing correction (Benjamini-Hochberg) was applied separately for each event type and data modality.

Gene-level mutation calls were obtained from Drews *et al*. [15]. For every gene and event type we calculated a Mann–Whitney U test between the event occurrences of samples that are and are not mutated in that gene and subsequently performed multiple-testing correction (Benjamini-Hochberg).

#### Modeling of internal event length distributions

For internal events, we fitted a shifted and scaled Pareto distribution of the form 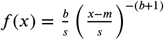 where *b* is the shape parameter, ***S*** the scale, and *m* the shift of the distribution, using the Pareto implementation in scikit-learn v1.7.0 [61]. Parameter estimates were *b* = 0.91, *m* = −1.20 × 10^6^, ***S*** = 1.20 × 10^6^ for gains, and *b* = 0.62, *m* = −4.87 × 10^5^, ***S*** = 4.88 × 10^5^ for losses. The required shift and scaling likely reflect the exclusion of segments shorter than 1 kbp, which truncates the distribution at the lower end.

#### Analysis of fragile site enrichment

Fragile sites were obtained from the HumCFS database [39]. Sites overlapping centromeres were excluded, yielding 108 fragile sites for analysis. Each site was padded by 50% of its original length, effectively doubling its size. We then counted the number of internal events overlapping each fragile site. To generate an expected background, we placed 10,000 random intervals per fragile site across the genome and counted overlapping internal events. The median count from these permutations was taken as the expected value under the null model of uniform event occurrence. Fragile sites were considered significantly enriched if the observed count exceeded 1.5 times the expected value.

#### The impact of whole-genome duplication on copy-number formation

Linear relationships were fitted using Random sample consensus (RANSAC), which robustly estimates linear trends by excluding outliers from the fit, as implemented in scikitlearn v1.7.0 [61].

### Generative model and detection of loci under selection

#### Event stratification and signal construction

Somatic copy-number events were stratified per chromosome into eight tracks defined by copy-number change (gain or loss) and four event-length classes as before while additionally limiting events to the range of 0.1Mbp-40Mbp to reduce noise: small (0.1–1 Mbp), mid1 (1–2.5 Mbp), mid2 (2.5–10 Mbp), and large (10–40 Mbp). For each track, internal events were aggregated into non-overlapping genomic bins to obtain event-count signals, using bin sizes matched to the characteristic scale of each class (4 kbp for small, 20 kbp for mid1, 40 kbp for mid2, and 200 kbp for large events). To assess variability in the observed signal, we generated 1,000 bootstrap replicates. Events were resampled 1,000 times per chromosome and class, aggregated into bins, and used to define 95% confidence intervals (2.5th–97.5th percentiles)

#### Generative model of SCNA formation

We developed a generative model of somatic copy-number alterations (SCNAs) based on the assumption that the observed event landscape arises from a combination of how events are generated and how they are selected. Event generation is modelled by uniform breakpoint formation along each chromosome, with the number and length distribution of events fixed to match the observed data. Selection is modelled as focal: if an event overlaps a locus of selection (oncogene-like or tumour-suppressor-like), it confers an evolutionary advantage (or disadvantage), increasing (or decreasing) the probability that such events are retained and thereby enriching events around these loci. In addition, we include a baseline component corresponding to events not influenced by any selected locus.

The generative model was implemented per chromosome and operates on internal SCNAs only, for which the assumption of uniform breakpoint formation is most appropriate. As input, it takes (i) the observed set of internal SCNA events for that chromosome, stratified into eight tracks, and (ii) a set of candidate loci under selection. Each locus is defined by its genomic position and eight track-specific selection strengths (nine parameters in total), with oncogene-like loci showing positive selection for gains and negative for losses, and tumour-suppressor-like loci exhibiting the opposite pattern.

In practice, for each track, each observed event is resampled along the chromosome by proposing *N*_steps_ random valid positions, where validity requires the event to remain internal and not overlap telomeres or the centromere. For a proposed position *p*, the weight was computed as

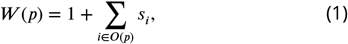

where ***O***(*p*) is the set of loci overlapping position *p*, and ***S***_*i*_ is the selection strength of locus *i* for the corresponding track. The baseline term (1) represents uniformly distributed neutral events, while positive or negative ***S***_*i*_ increase or decrease the likelihood of retaining overlapping events. Selection strengths are in arbitrary units, defined relative to each other and to the neutral baseline of 1. For each event, the weights over all proposed positions are normalized to obtain a probability distribution along the chromosome, which is then mapped to the binning scheme of the corresponding track. Longer events contribute to more bins and thus have proportionally larger influence on the simulated signal. Summing these position-wise probabilities over all events in the track yields the expected SCNA occurrence profile for that track.

See also 1 for a pseudo-code description of the generative model.

##### Algorithm 1

Generative model simulation of SCNA event profiles

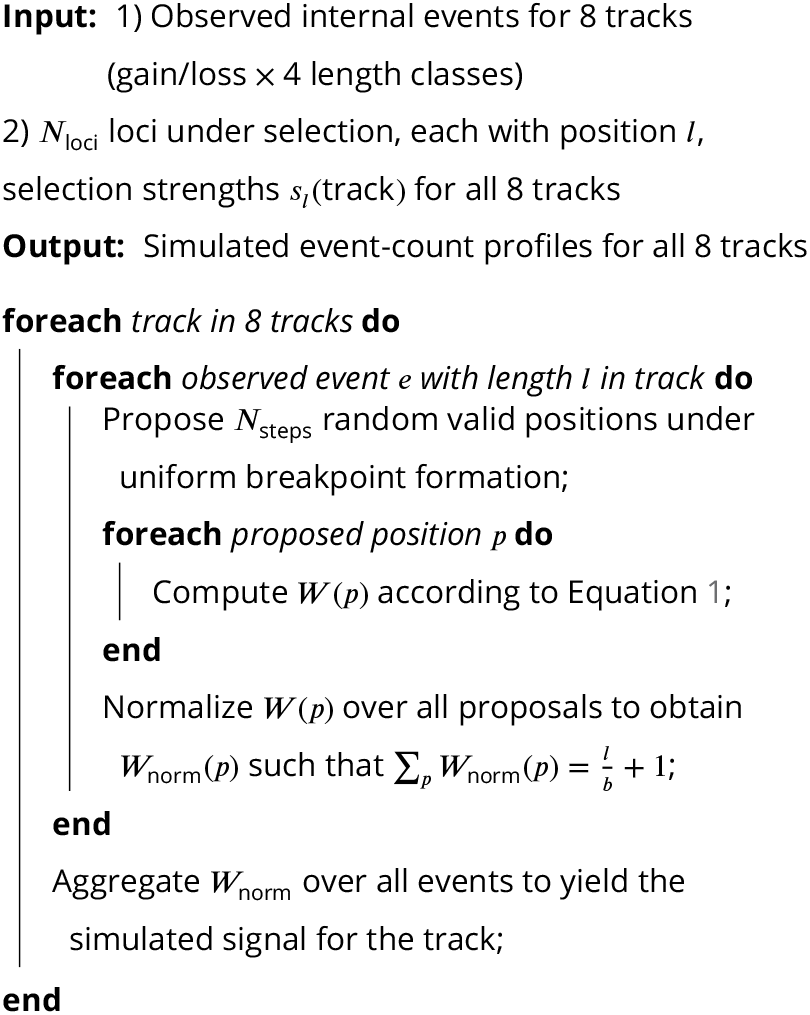

#### Convolution approximation to the simulation

Because explicit event-level simulation was computationally intensive, we implemented an equivalent convolution-based approximation. First, a kernel was generated by simulating a single locus of selection with selection strength ***S***_*i*_ = 1 in the absence of a baseline rate, producing a template for the characteristic triangular shape. A selection track was then constructed as an array matching the genomic bins of the observed signal, set to 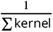 everywhere and to 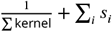 at bins containing selected loci. Convolving the kernel with this selection track yielded the simulated signal, effectively placing and scaling the triangular enrichments on top of the uniform baseline. A neutral model without loci was simulated, normalized to maximum value 1, and used to scale the convolution output near telomeres and the centromere. The output of the convolution simulation was then scaled such that the area under the curve for the convolution simulation matched the area of the real data. This approach produced results nearly identical to the full simulation while being substantially more efficient.

#### Detection of loci under selection

Loci under selection were inferred using an iterative Markov Chain Monte Carlo (MCMC) approach that jointly optimizes the positions and selection strengths of all loci across the eight event tracks. The procedure starts without any loci and iteratively adds new loci at regions showing the largest discrepancy between observed and simulated signals. After each addition, the positions and selection strengths of all existing loci are reoptimized simultaneously to improve the overall model fit. To prevent loci from being placed too close together, a 20 kbp region around each locus is blocked from further placement.

Optimization used a simulated annealing scheme [62]. At each iteration, one locus was randomly chosen and modi-fied—either a position shift (10% of steps, uniformly within ±3 × 10^5^ bp) or a change to one of its selection strengths (90%, drawn uniformly between 0 and the current maximum strength). The modified locus set was evaluated by generating simulated event tracks and computing the mean squared error (MSE) against the observed signal. Proposals were accepted if they improved the fit or, otherwise, with probability

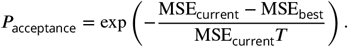

The temperature ***T*** followed the schedule ***T***_*i*_ = 10^−3−2*i*/*n*^, for iteration *i* and total number of iterations *n*. High temperatures early in the process allow acceptance of suboptimal moves to escape local minima, while later cooling favors only improvements, guiding convergence toward an optimal solution.

For each chromosome, 100 loci were placed and subsequently refined through post-processing and p-value filtering (see below). Positional uncertainty was estimated using 1,000 bootstrap replicates, re-optimizing each locus position per replicate while keeping others fixed; bounds were defined as ±1 s.d. around the original locus position (total width = 2 s.d.), with a minimum bound set to 1/4 of the kernel width of the smallest length scale to account for the uncertainty of the triangular shape.

#### Post-processing of loci

Nearby loci of the same type (OG–OG or TSG–TSG) with overlapping bounds were merged. To simplify the model, selection strengths were set to zero if removal did not push simulated signals outside the 95% confidence interval. Loci were further filtered if they had positional uncertainty exceeding the mean length of the largest active track, or a triangular prominence < 10 in all tracks, where prominence denotes peak height relative to the lowest surrounding minimum.

To assess statistical support for inferred loci, we computed empirical p-values using a permutation test based on the same neutral simulation used in the generative model, i.e. event shuffling under uniform breakpoint formation with no loci of selection. For each chromosome and polarity (oncogene-like for gains; tumour-suppressor-like for losses), internal events were shuffled to random valid positions along the chromosome. In each neutral replicate, we selected a random genomic position and fit a single fixed-position locus using the same MSE-based optimization procedure as for locus detection, and assigned events to this locus and the baseline component using the same probabilistic assignment scheme as for observed loci (see below). The test statistic was the expected number of events assigned to the locus, ***S*** = ∑_*e*_*P*_*i*_ (*e*), summed across length scales. Repeating this procedure 10,000 times yielded a null distribution of ***S*** under neutrality. For each inferred locus, the empirical p-value was the fraction of null replicates with ***S***_null_ ≥ ***S***_obs_. P-values were adjusted genome-wide using Benjamini-Hochberg, and loci with adjusted p-values *<* 0.05 were retained.

Genes were assigned to loci based on genomic overlap: any gene whose coordinates fell within the positional bounds of a locus was linked to that locus. For naming purposes in the main text, each locus was represented by a single gene, prioritizing known oncogenes or tumour-suppressor genes from the COSMIC database [40]. If no COSMIC gene was present, the top candidate was selected from the Davoli gene list [37]. Although a locus may contain multiple genes, only the highest-priority gene was used for display and reference.

### Assigning observed events to individual loci and calculating the baseline rate

Using the inferred locus positions and selection strengths, each observed event was probabilistically assigned to one or more loci, or to the baseline component. For a given evente, we first identified the set of overlapping loci O(e) on the same chromosome. Events with no overlaps were assigned entirely to the baseline. For overlapping events, we computed assignment probabilities rather than hard labels, reflecting the relative contribution of each locus. For N loci, the resulting assignment vector has length N+1 (the additional element representing the baseline rate).

The probability that event *e* belongs to a specific overlapping locus *i* ∈ *O*(*e*) was defined as 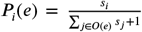, where ***S***_*i*_ is the track-specific selection strength of locus *i*. The corresponding baseline probability was 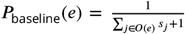. The total number of events per locus and the overall number of baseline events were obtained by summing *P*_*i*_(*e*) and *P*_baseline_(*e*) across all events, respectively.

To assess whether baseline rates reflected true genomic patterns rather than artifacts of event inference, we quantified breakpoint uniformity in the raw copy-number profiles. Break-point occurrences were counted in 200 equally sized bins per chromosome, excluding centromeric regions. For each chromosome, we calculated the median breakpoint count per bin and the mean absolute deviation (MAD) from this median. Under uniform breakpoint formation, the MAD is low, producing a flat histogram, whereas clustering near loci under selection results in higher deviation. The MAD per chromosome was then correlated with the inferred baseline rate to test whether higher baseline rates corresponded to more uniform break-point distributions.

### Comparison to previously reported loci and enrichment analyses

Loci reported by GISTIC [6] and BISCUT [22] were used as benchmarks, as both were calculated on TCGA data. Loci were defined using each method’s multiple-testing–adjusted significance threshold at *p* < 0.05. GISTIC and BISCUT loci were subsequently filtered to remove large loci (>50 kbp) and loci located inside of centromeres. For BISCUT, both positively and negatively selected loci were considered. GISTIC loci were padded to a minimum width of 25 kbp to facilitate overlap calculations. We excluded cancer type-specific GISTIC loci because, even after removing loci >10 Mbp, their union spans 24% of the genome and most occur only in a small subset of cancer types, making them not directly comparable to our pan-cancer analysis.

Loci identified by SPICE were compared to these previous calls by defining overlaps where start and end coordinates intersected. For three-way comparisons among SPICE, GISTIC, and BISCUT, GISTIC and BISCUT loci were first overlapped to form pairs, and SPICE loci were considered overlapping if they fell within the union of these paired intervals. Remaining SPICE loci were then compared to unmatched GISTIC or BISCUT loci.

Significance of overlap was assessed using a permutation test. For each SPICE locus, 10,000 valid genomic positions were randomly sampled (excluding centromeres), and the number of overlaps with GISTIC, BISCUT, or COSMIC genes was calculated for each permutation. P-values were defined as the fraction of permutations with equal or greater overlap than observed. COSMIC gene lists were downloaded from the official COSMIC database (accessed Feb 26, 2025) [40]. This process was repeated with the mutation-based TSG/OG list published by Davoli *et al*. [37].

## Availability of data and materials

SPICE is implemented in Python 3 and freely available under GPLv3 on GitHub: https://github.com/ICCB-Cologne/spice. Inferred loci are available under https://doi.org/10.5281/zenodo. 18751184.

## Acknowledgements

TLK kindly thanks Klaus-Robert Müller for support. Computation has been performed on the HPC for Research cluster of the Berlin Institute of Health. The results shown here are in part based upon data generated by the TCGA Research Network: https://www.cancer.gov/tcga. The authors furthermore thank Katyayni Ganesan and Chenxi Nie for feedback on the manuscript.

## Author contributions

TLK and RFS designed the method. TLK implemented the method and processed and analysed the data. TKL, AS, FM, PVL, RFS interpreted the results. TLK, RFS, FM and PVL wrote and edited the manuscript. AS derived the mathematical proof for the event reconstruction. All authors read and approved the final manuscript version.

## Supplementary figures

**Supplementary Figure 1.**
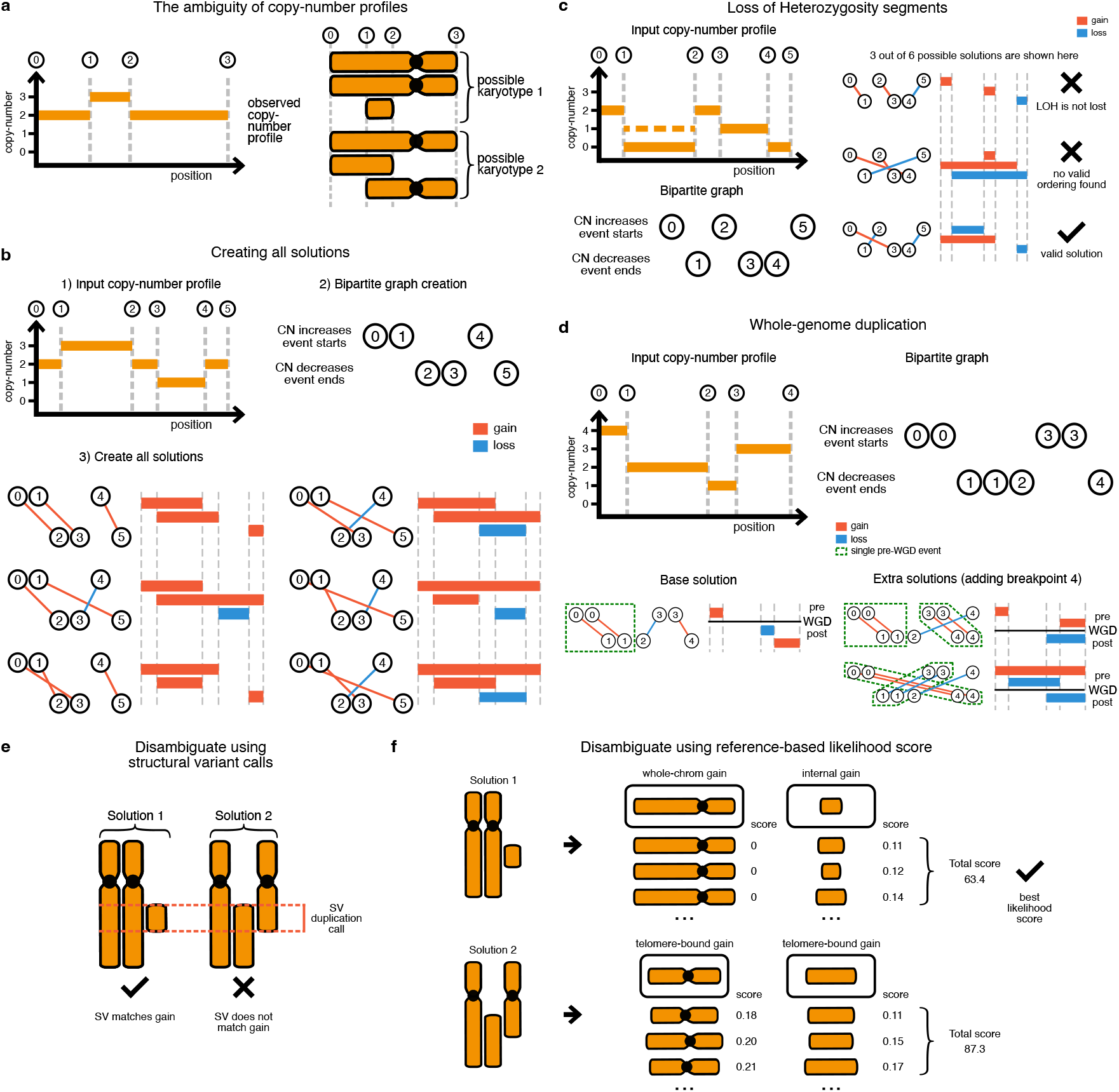
Event inference framework and bipartite graph representation. **a**, Copy-number profiles are ambiguous with respect to the underlying karyotype. **b**, Example of the event inference framework. Copy-number profiles are converted into bipartite graphs where each node corresponds to a breakpoint. All solutions to the bipartite graph are then converted into copy-number events (gains and losses) where each edge in the graph corresponds to a single event. **c**, Illustration of loss-of-heterozygosity (LOH) segments, showing invalid and valid event orderings consistent with full allele loss. **d**, Representation of pre- and post–whole-genome duplication (WGD) events, showing pre-WGD events as duplicated edges. **e**, Selection of multiple solutions using structural variant (SV) calls. **f**, Selection of the most plausible solution by similarity scoring against the PCAWG reference set, integrating structural variant (SV) support when available.

**Supplementary Figure 2.**
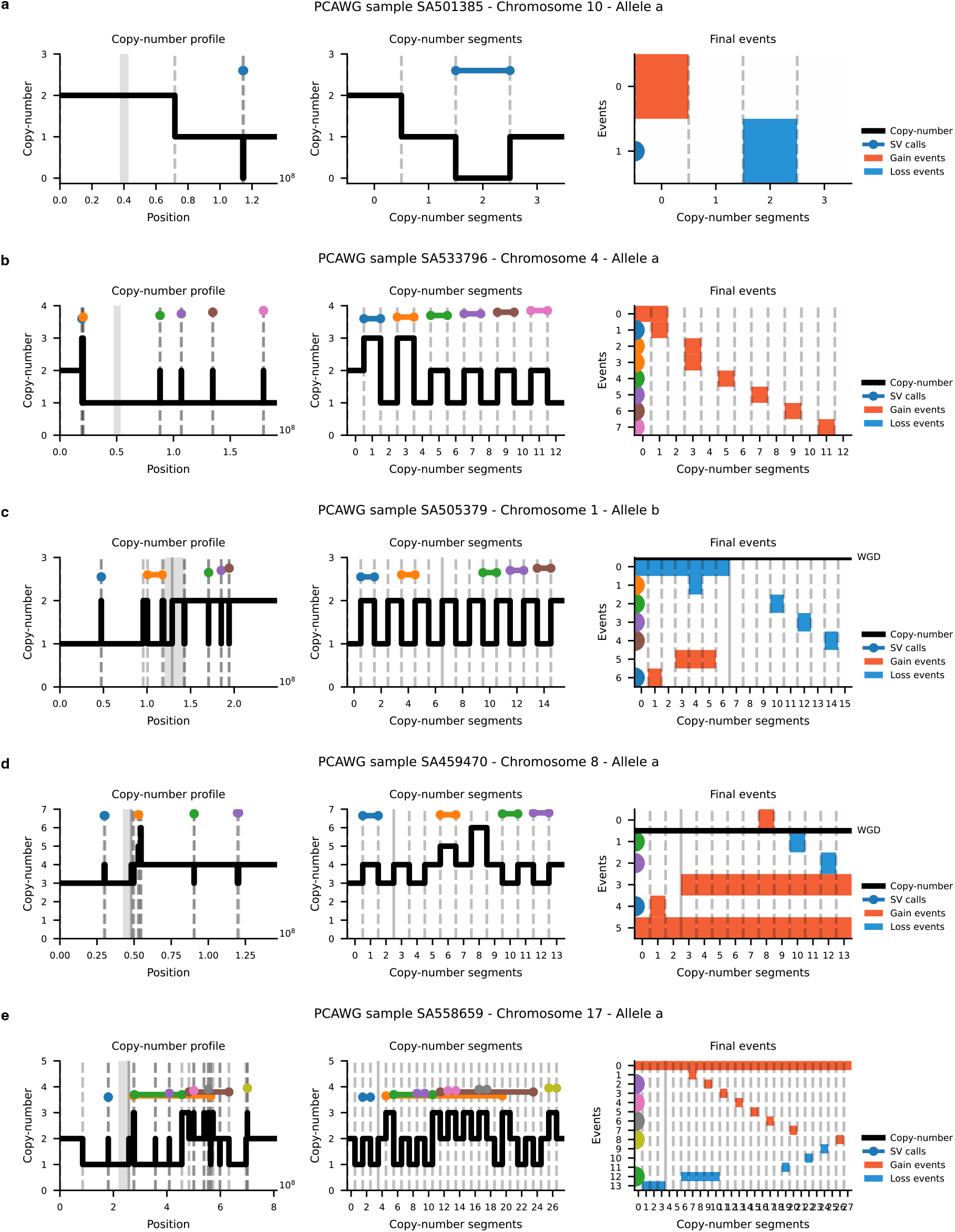
Examples of inferred events. Representative examples of allele-specific copy-number profiles, corresponding copy-number segments, and final inferred events for selected PCAWG samples. Each panel shows (left) the raw copy-number profile, (middle) segmented copy-number representation, and (right) the inferred gain (red) and loss (blue) events after applying the minimum-evolution framework. Structural variant (SV) calls are shown for validation.

**Supplementary Figure 3.**
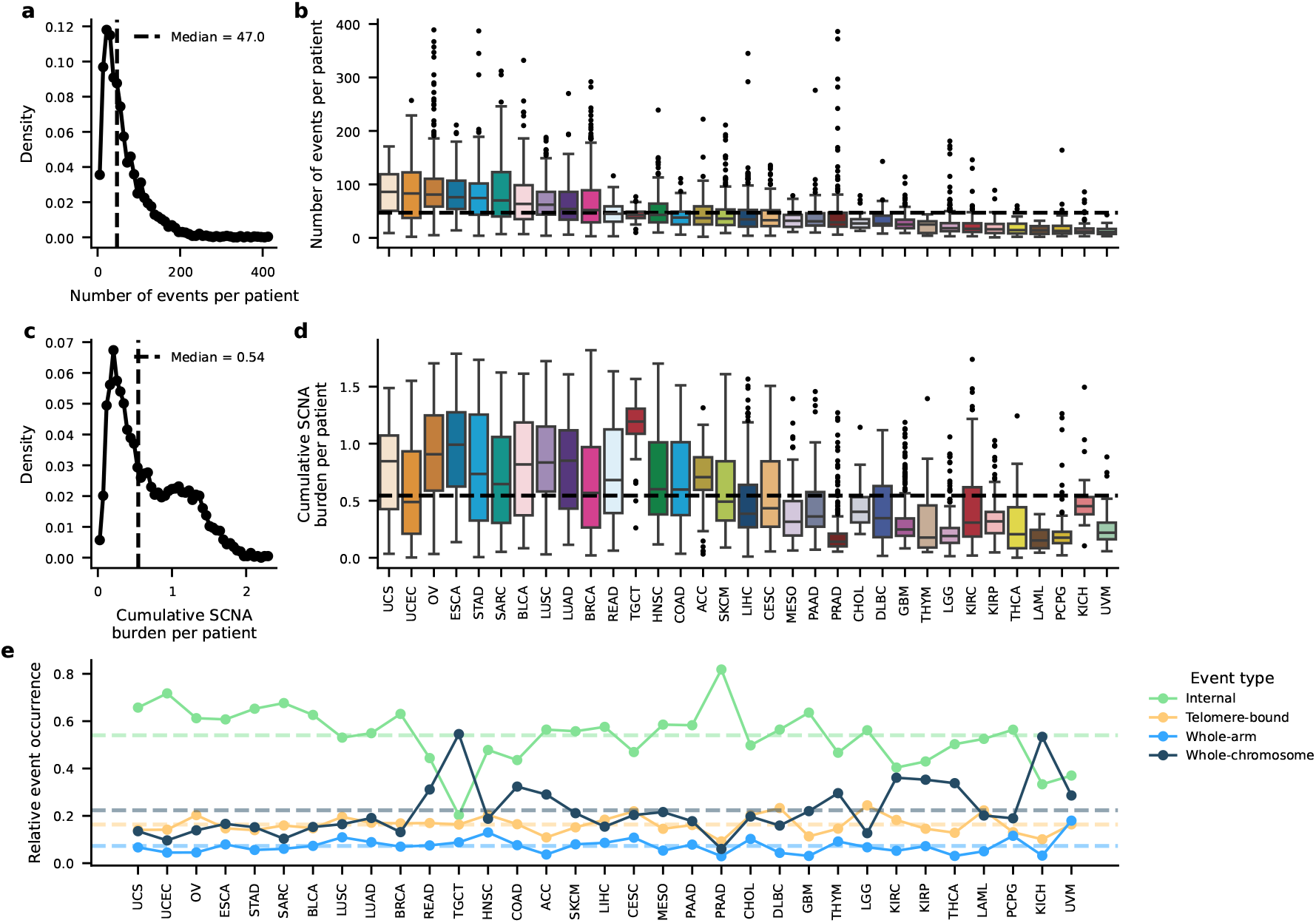
Genome-wide distributions of inferred copy-number events across 5,966 TCGA samples. **a**, Distribution of the total number of somatic copy-number alteration (SCNA) events per patient. **b**, Number of SCNA events per patient across 33 TCGA cancer types. **c**, Distribution of the cumulative SCNA burden per patient calculated as the sum of inferred event lengths divided by the haploid genome length. As overlapping events are counted multiple times values can exceed 1.0. **d**, Cumulative SCNA burden per patient, stratified by cancer type. **e**, Relative occurrence of event types (internal, telomere-bound, whole-arm, and whole-chromosome) across cancer types.

**Supplementary Figure 4.**
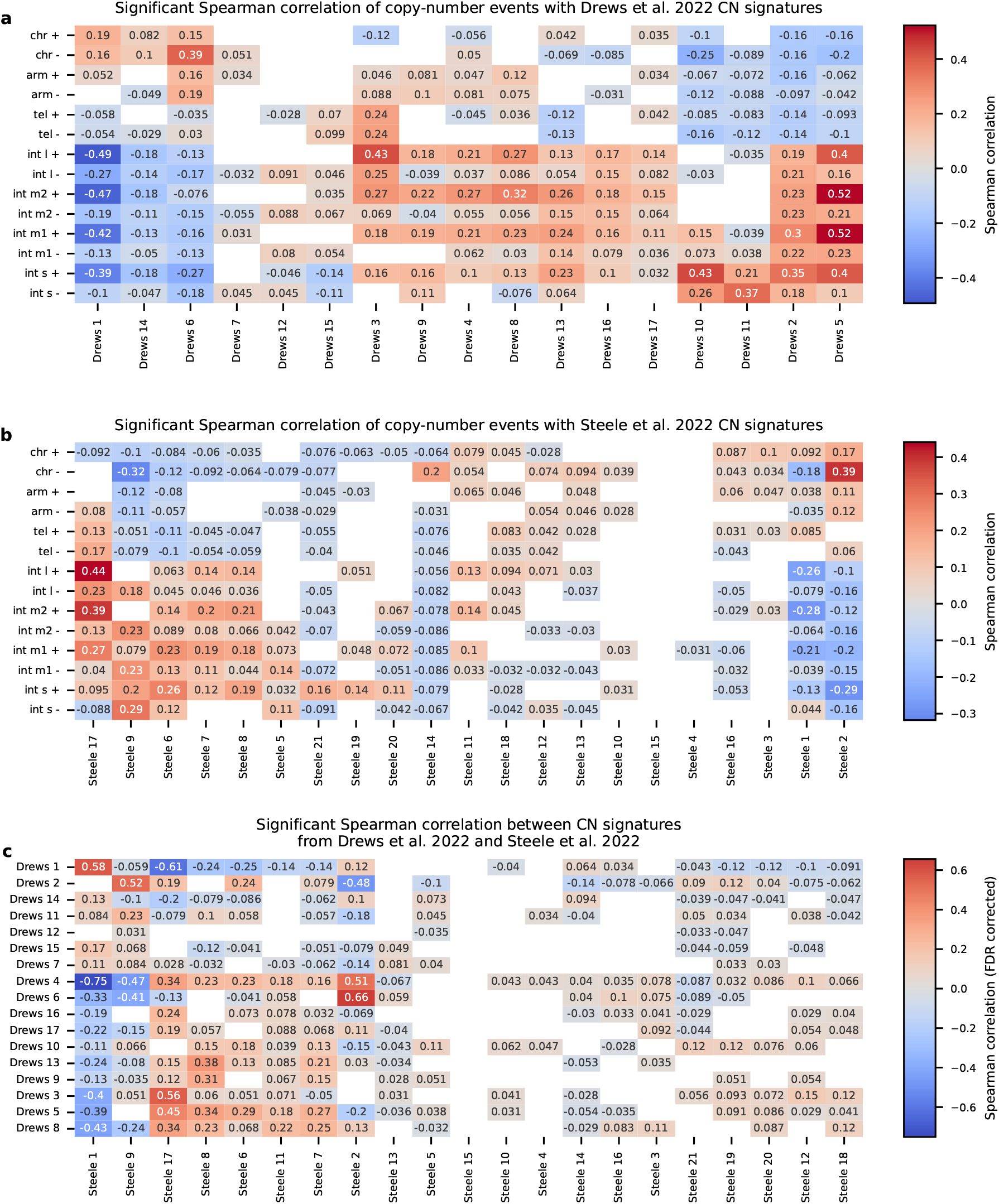
Correlation of inferred copy-number event types with published copy-number (CN) signatures. **a**,b Spearman correlations between the per-patient frequencies of SCNA event types and copy number signatures from Drews et al. 2022 (a) as well as Steele et al. 2022 (b). **c**, Correlation matrix comparing copy number signatures from Drews et al. 2022 and Steele et al. 2022, showing limited direct correspondence between the two frameworks. All correlations are Spearman rank coefficients; colour scale denotes correlation strength, and only FDR-corrected significant values are displayed.

**Supplementary Figure 5.**
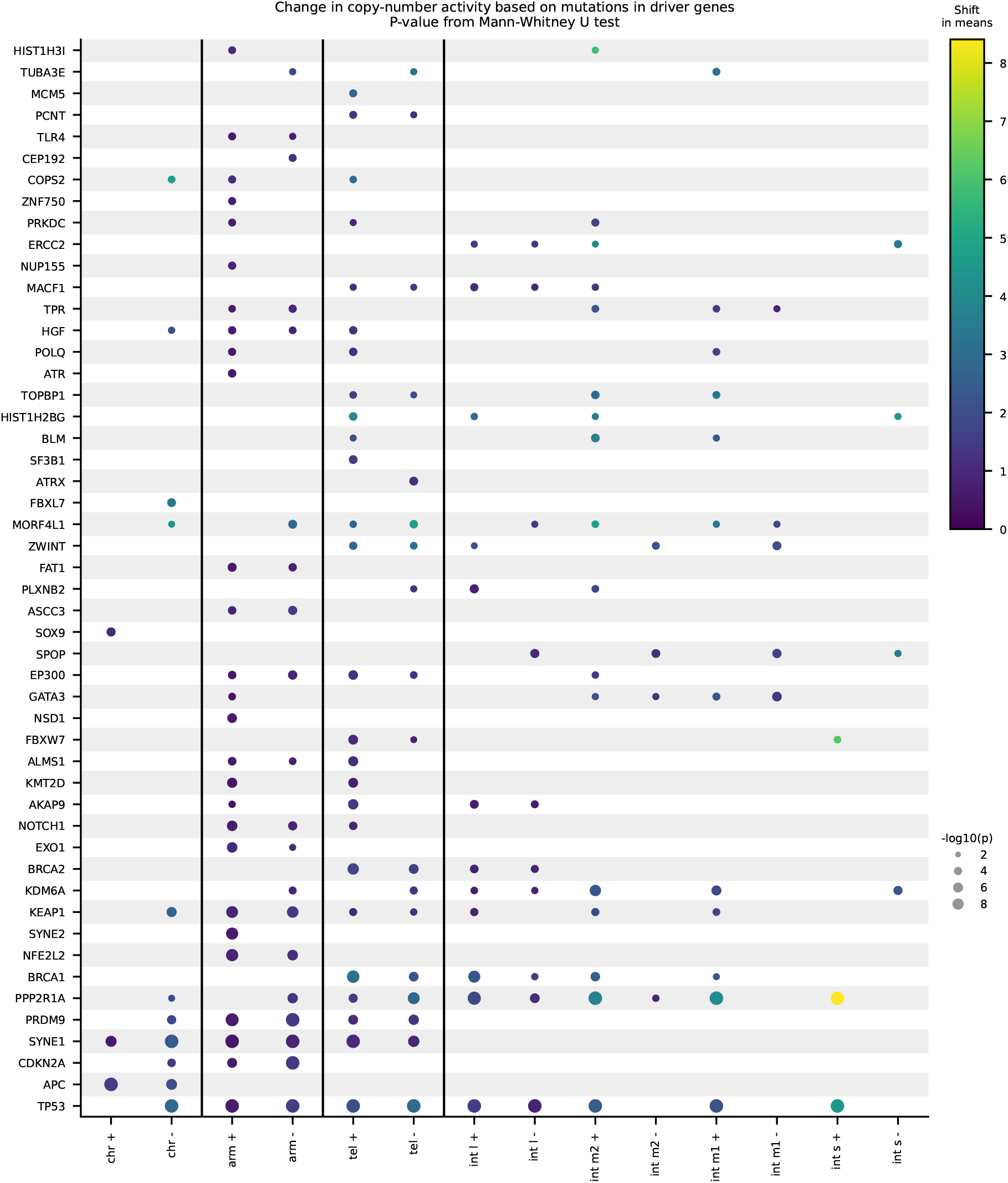
Associations between driver gene mutations and somatic copy-number alteration (SCNA) activity. Each dot represents a significant association between mutations in a driver gene (rows) and the number of copy-number events of a given type (columns), computed using a Mann–Whitney U test. Dot size encodes significance (− log_10_(*P*)), and colour represents the shift in means between mutated and non-mutated samples.

**Supplementary Figure 6.**
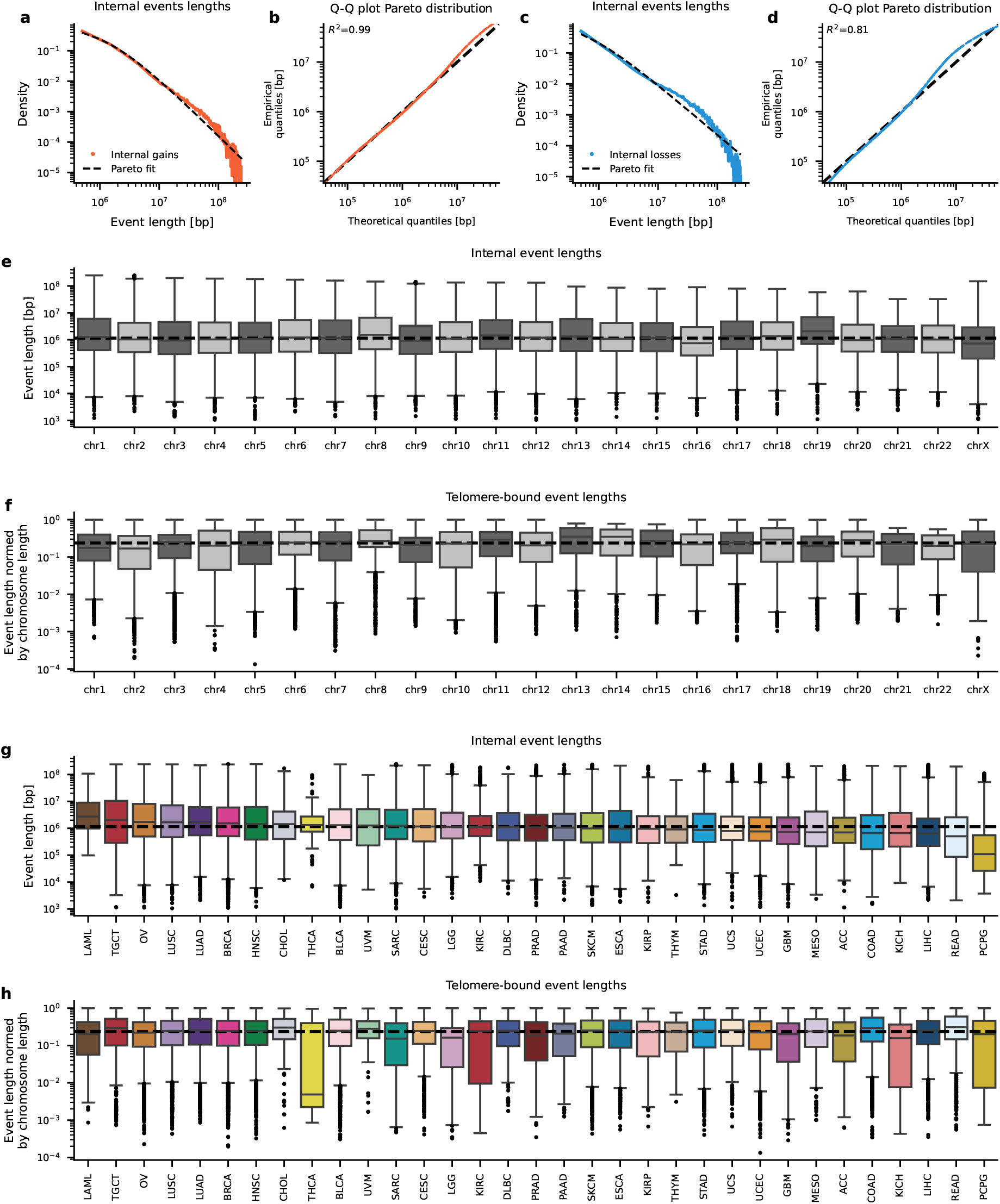
Length distributions of internal and telomere-bound somatic copy-number events. a–d, Internal event length distributions and Q-Q plots for gains (a,b) and losses (c,d), fitted with Pareto distributions (***r***^2^gain = 0.99, ***r***^2^loss = 0.81). **e−f**, Event lengths per chromosome for internal (e) and telomere-bound (f) events. **g−h**, Event lengths per cancer type for internal (g) and telomere-bound (h) events, showing broad consistency across chromosomes and tumour types.

**Supplementary Figure 7.**
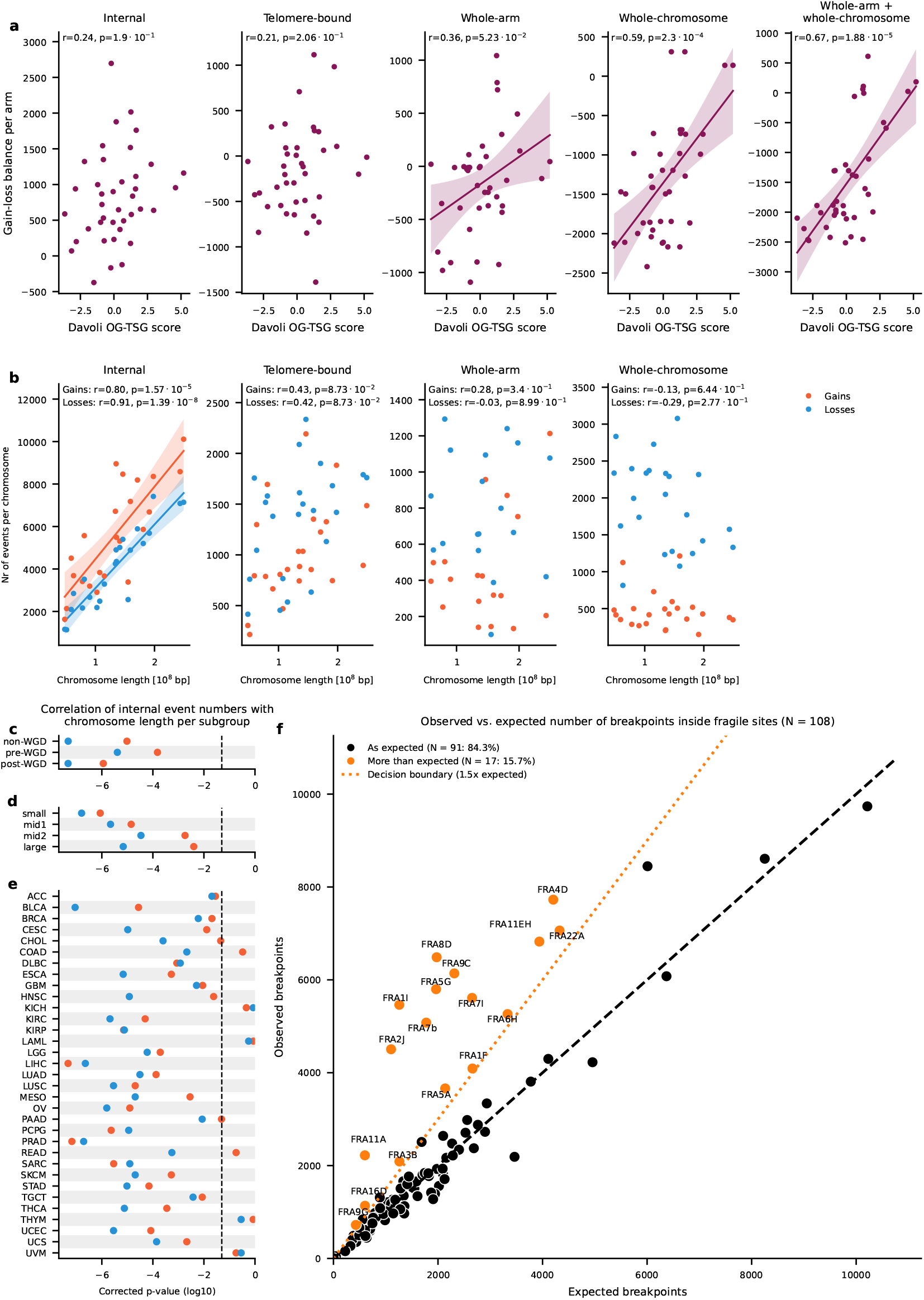
Chromosome-level selection patterns and fragile site analysis. **a**, Correlation between gain–loss balance of copy-number events and oncogene–tumour suppressor (OG–TSG) scores from Davoli et al. [37] per chromosome arm. **b**, Relationship between chromosome length and number of events per chromosome. **c–e**, Correlation coefficients (***r***) between internal event counts and chromosome length, stratified by WGD status (c), internal event length class (d) and cancer type (e). **f**, Observed versus expected number of breakpoints within common fragile sites (HumCFS database).

**Supplementary Figure 8.**
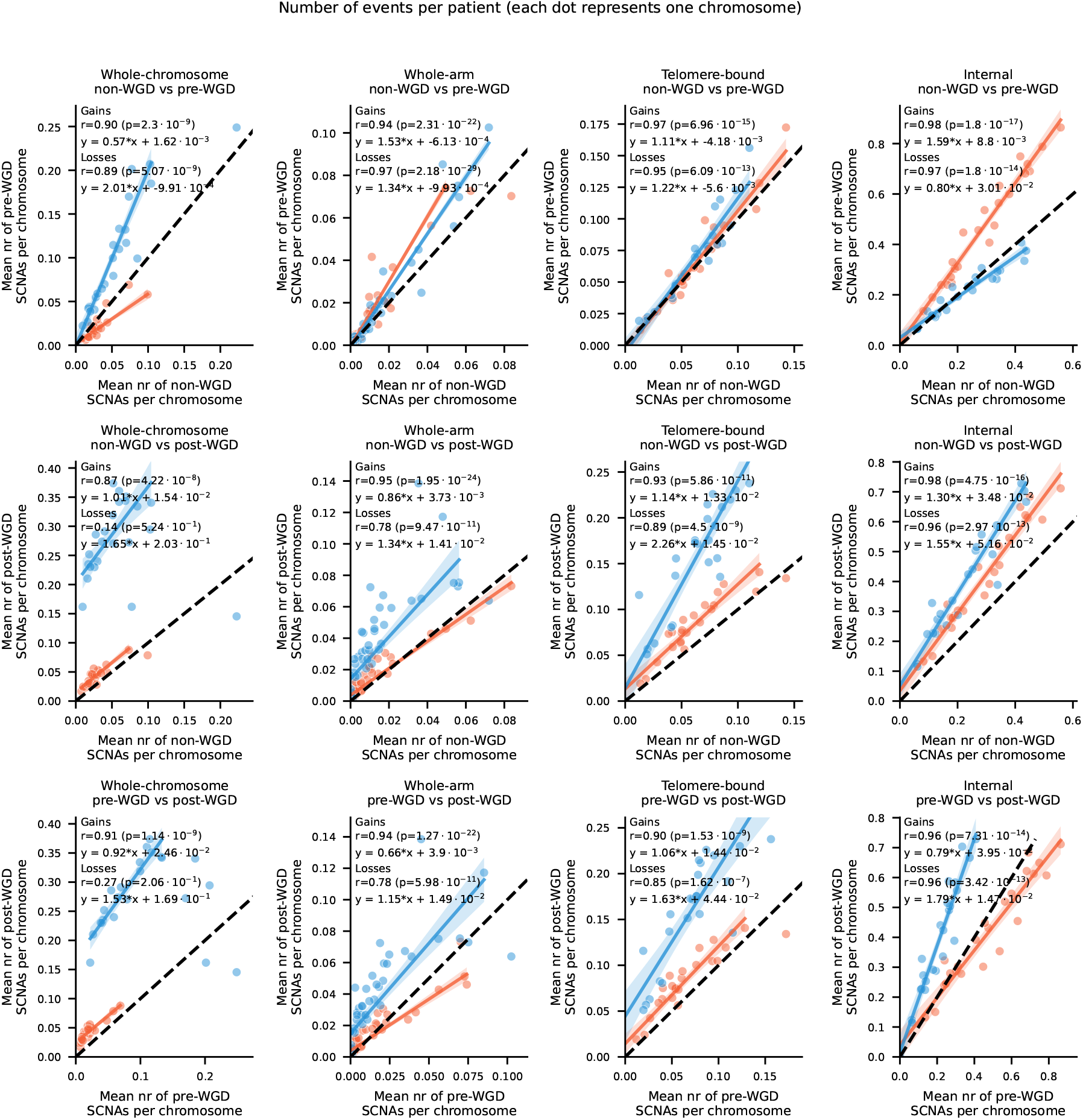
Comparison of somatic copy-number alteration (SCNA) event frequencies across whole-genome duplication (WGD) states. Each dot represents one chromosome. Linear relationships were fitted using random sample consensus fitting.

**Supplementary Figure 9.**
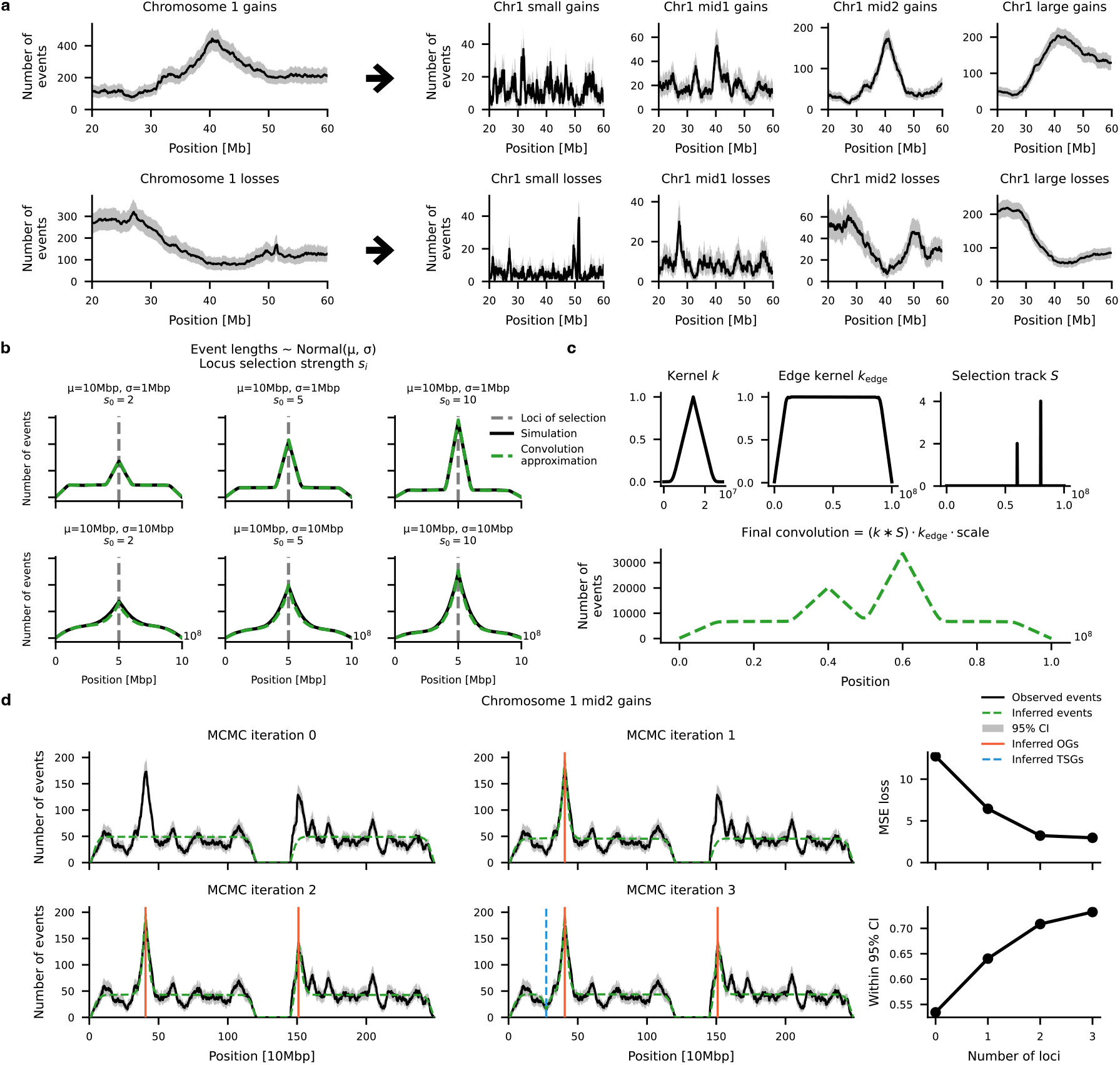
Generative model and convolution-based approximation for detecting loci under selection. **a**, Example of event stratification by length scale for chromosome 1, showing separation of gains and losses into four classes (small, mid1, mid2, large). **b**, Simulated enrichment shapes generated for varying locus selection strengths (***S***_*i*_) and event length distributions (*μ, σ*). Increasing ***S***_*i*_ enhances the amplitude of the triangular signal, while wider length distributions broaden its base. **c**, Convolution-based approximation of the generative model. A triangular loci kernel (*k*) convoluted with the selection track (***S***) and finally multiplied by an edge kernel (*k*_edge_) to generate the expected event profile. **d**, Iterative Markov Chain Monte Carlo (MCMC) inference procedure for chromosome 1 mid2 gains. Successive iterations add or adjust loci under selection (red = oncogene-like, blue = tumour-suppressor–like), progressively improving the fit of the model (green) to the data (black) and reducing mean squared error (MSE) while increasing the fraction of the genome within 95% confidence intervals.

**Supplementary Figure 10.**
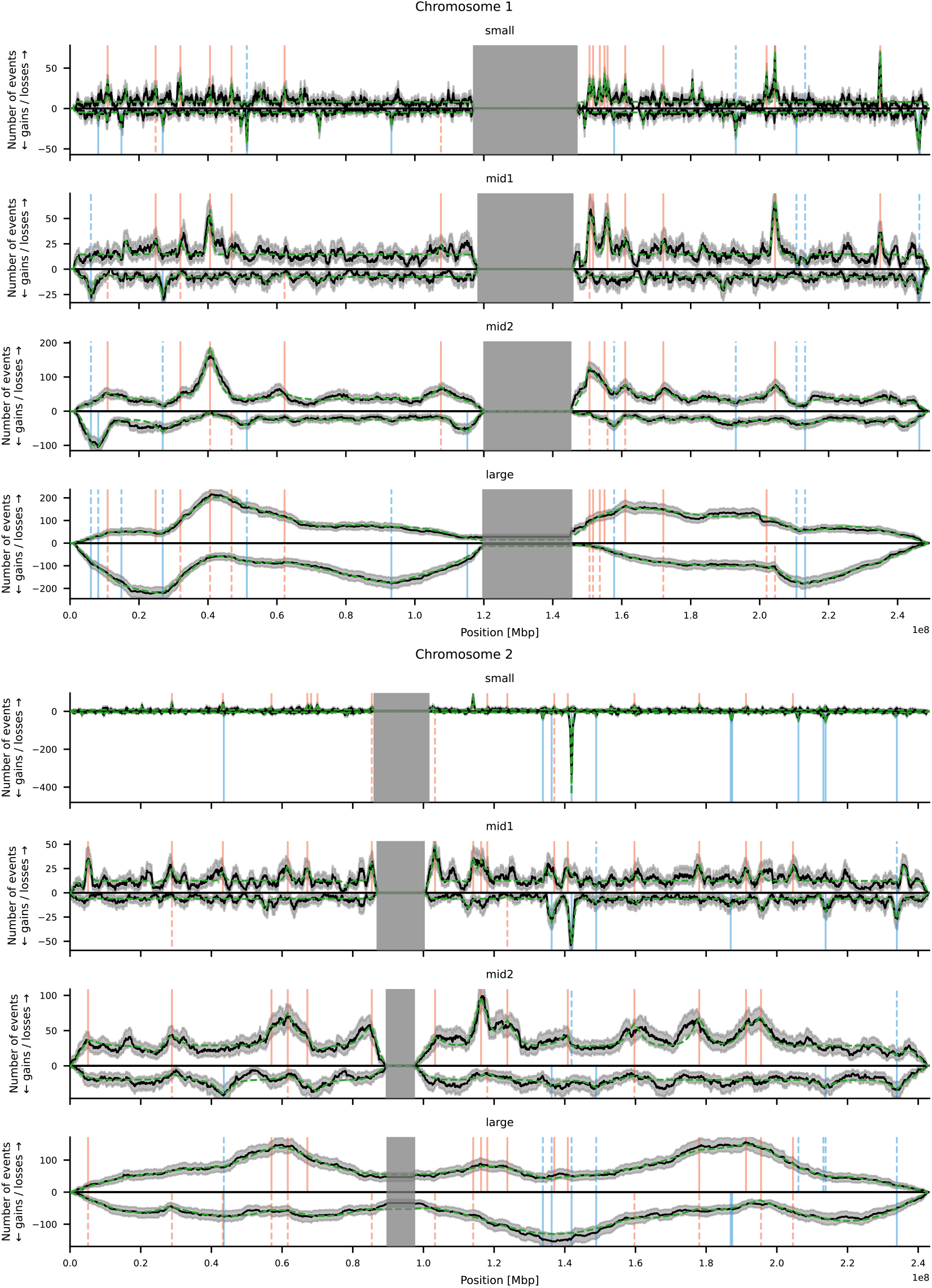
Generative model fit for chromosomes 1 and 2 across four event-length scales. Observed and simulated event-count profiles are shown for chromosome 1 (top) and chromosome 2 (bottom), stratified into four internal event-length classes: small (0.1–1 Mbp), mid1 (1–2.5 Mbp), mid2 (2.5–10 Mbp), and large (10–40 Mbp). Black lines represent observed event counts, with shaded regions indicating 95% bootstrap confidence intervals. Green lines denote model fits generated by the convolution-based approximation, while red and blue vertical lines mark inferred oncogene-like and tumour-suppressor–like loci, respectively. Grey boxes indicate centromeric regions excluded from analysis.

**Supplementary Figure 11.**
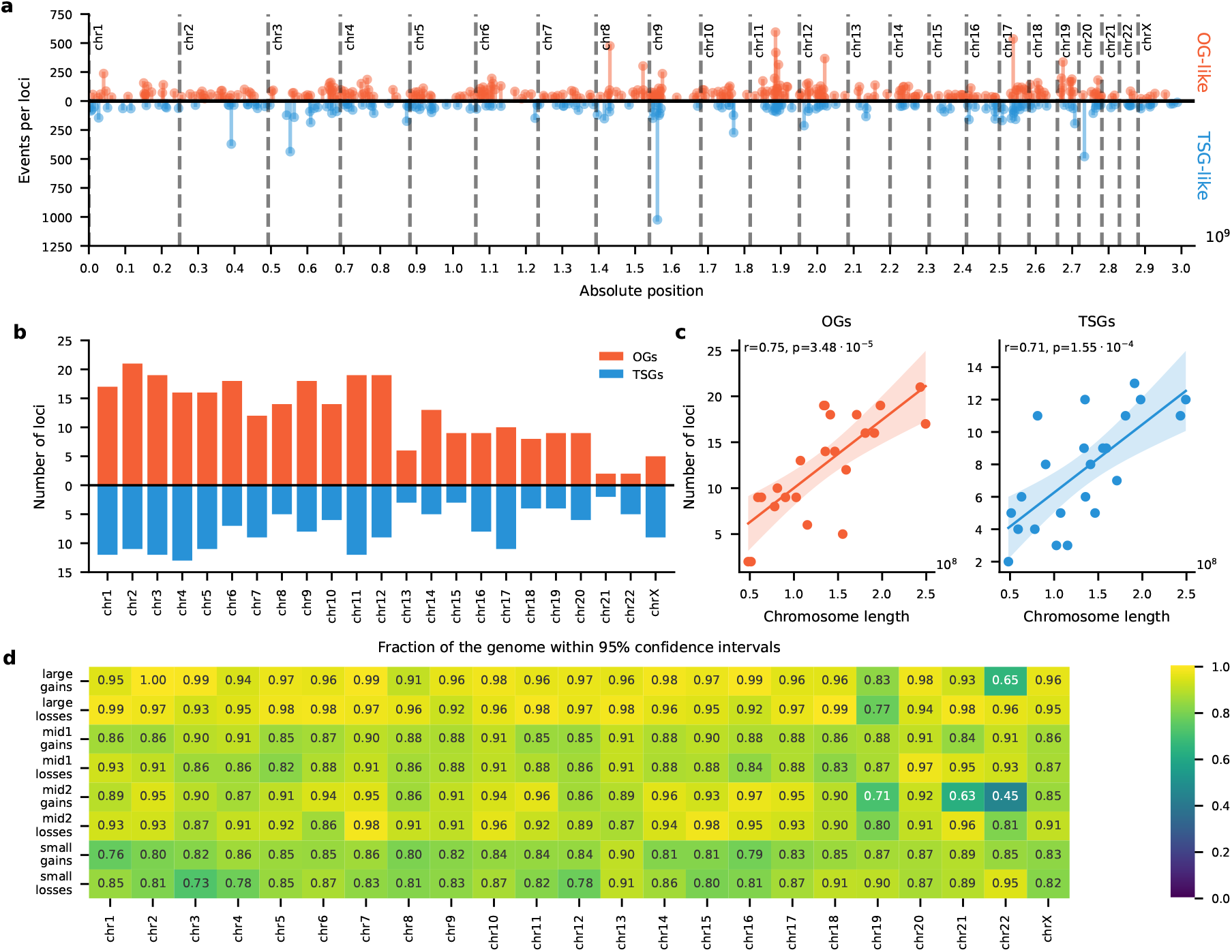
Genome-wide landscape of loci under selection. **a**, Distribution of inferred loci across the genome, showing oncogene-like (OG-like, red) and tumour-suppressor–like (TSG-like, blue) selection signals. Several well-established cancer genes (e.g., CDKN2A, FHIT, SOX17, RBFOX1) appear among top loci. **b**, Counts of inferred loci per chromosome. **c**, Correlation between the number of inferred loci and chromosome length for OG-like (left) and TSG-like (right) loci. **d**, Fraction of the genome with generative model within the 95% confidence intervals of the observed event counts, stratified by chromosome, event class and gain/loss.

**Supplementary Figure 12.**
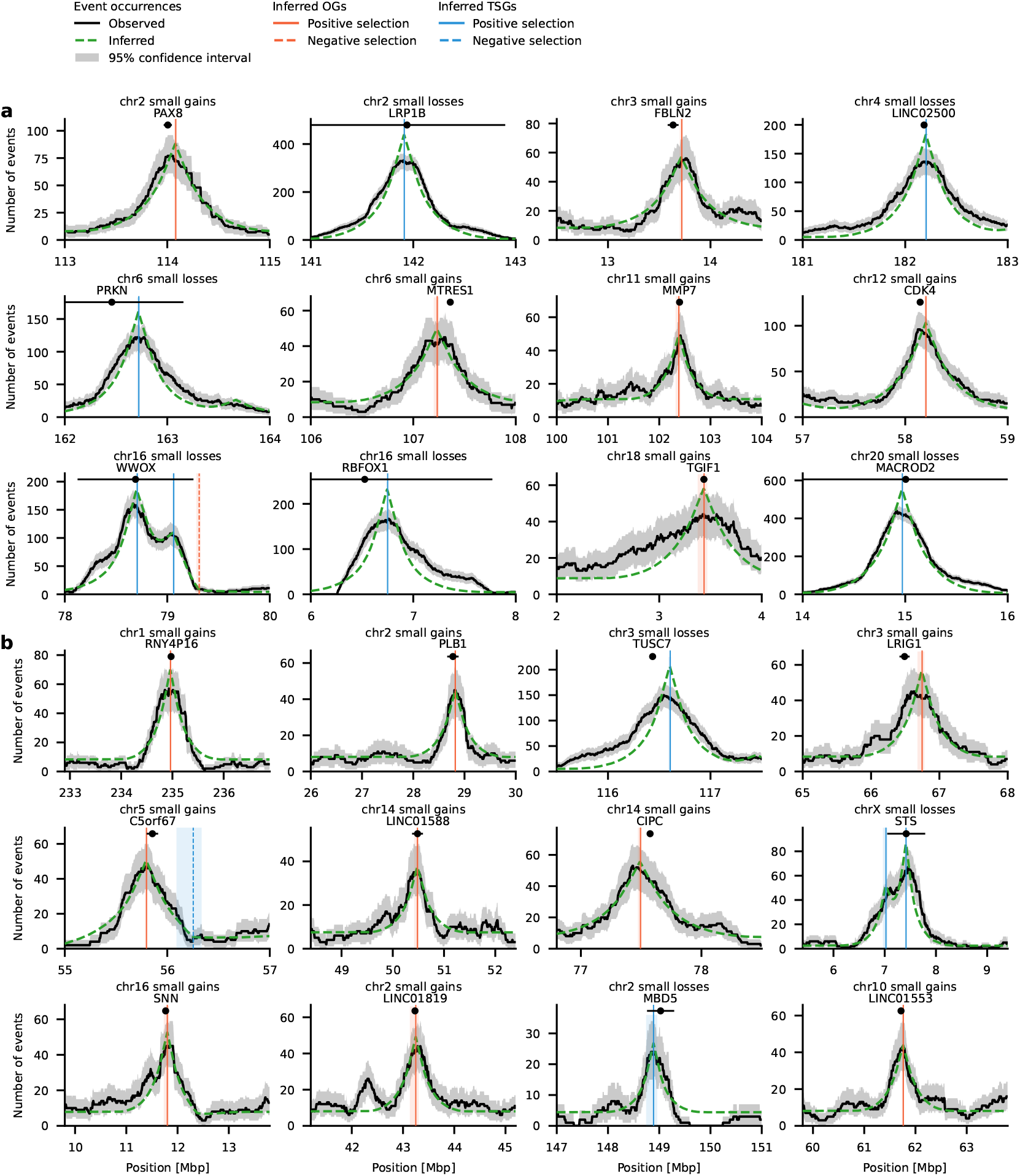
Example loci identified by SPICE corresponding to known and novel regions under selection. Event occurrence profiles across small event length scales for representative loci. a, Examples of loci previously reported by GISTIC or BISCUT. **b**, Examples of novel loci detected by SPICE not present in prior analyses.

**Supplementary Figure 13.**
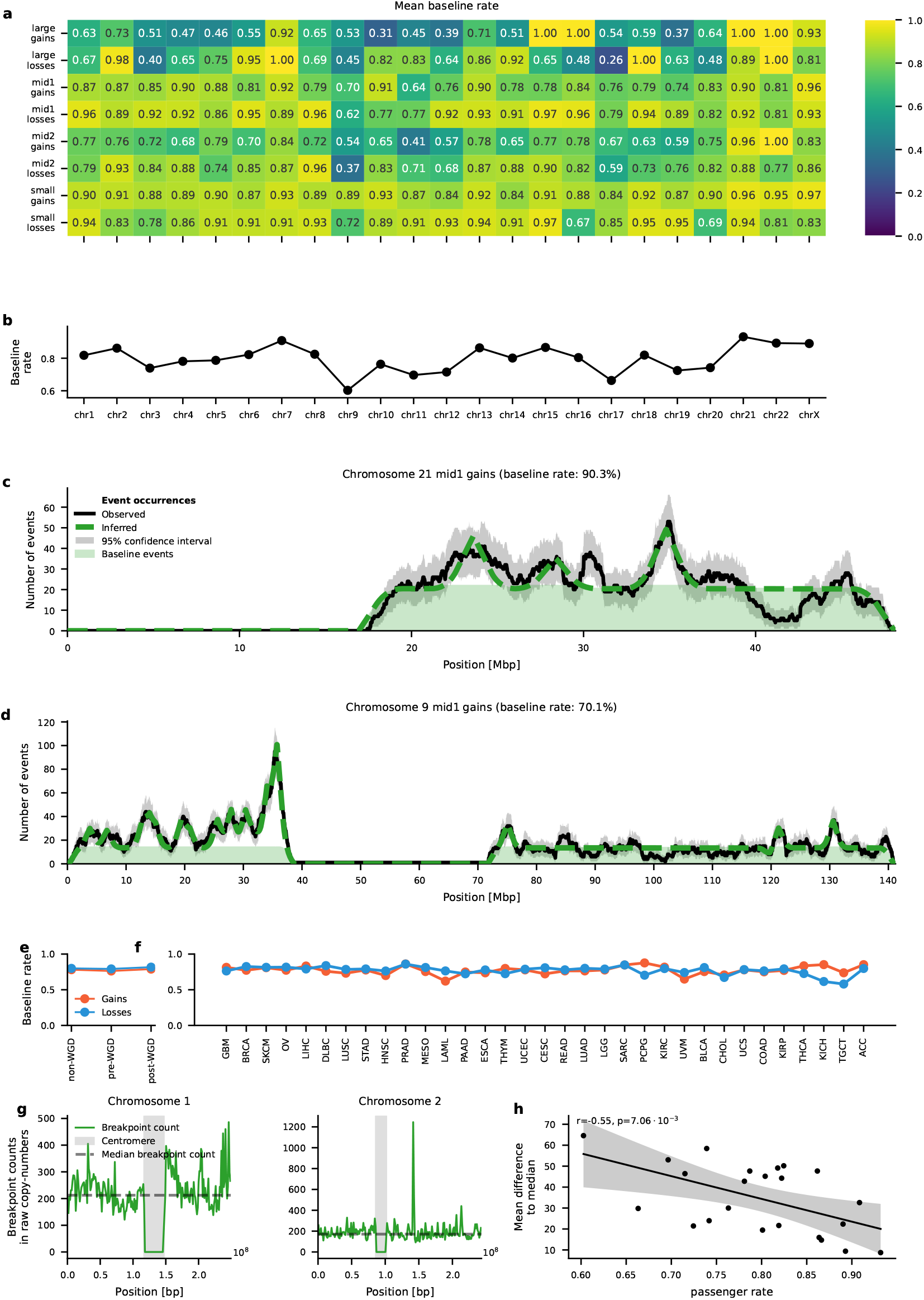
Baseline event rate estimation. **a**, Mean baseline event rates per chromosome and event class. **b**, Genome-wide distribution of mean baseline rates across chromosomes. **c–d,** Examples of chromosomes with contrasting baseline rates. For chromosome 21 SPICE slightly overestimates counts around 41 Mbp because a negatively selected locus did not pass the p=0.05 threshold. **e–f**, Baseline rates stratified by WGD status (e) and cancer type (f). **g**, Example breakpoint density profiles for chromosomes 1 and 2. h, Correlation between chromosome-wise baseline rate and breakpoint uniformity (MAD from the median).

**Supplementary Figure 14.**
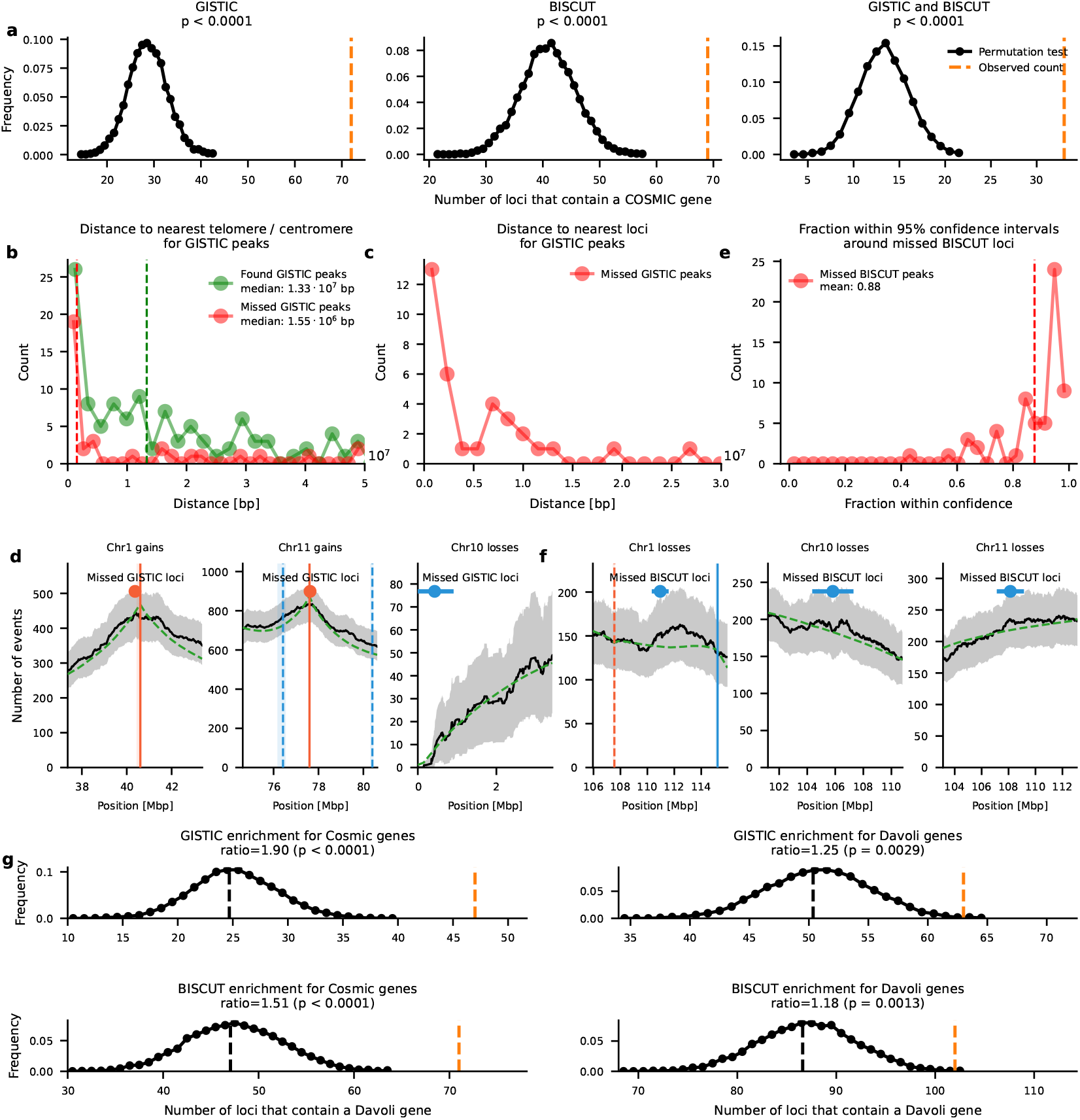
Comparison of SPICE loci with previously reported GISTIC and BISCUT calls. **a**, Permutation tests showing that the number of SPICE loci overlapping GISTIC, BISCUT, and joint GISTIC–BISCUT loci is significantly higher than expected by chance. **b**, Distances of matched (green) and missed (red) GISTIC loci to the nearest telomere or centromere, indicating that missed loci are frequently located near low-mappability regions. **c**, Distances of missed GISTIC loci to the nearest SPICE locus, showing close spatial proximity (<1 Mbp) for most. **d**, Examples of missed GISTIC loci at chromosomes 7, 10 and 11, illustrating local enrichment consistent with nearby SPICE peaks as well as proximity to the telomere. **e**, Distribution of model-data agreement (fraction of bins within 95% confidence intervals) for missed BISCUT loci, indicating high concordance (median = 0.84). **f**, Examples of missed BISCUT loci with model-data agreement in the absence of inferred loci. **g**, Enrichment analyses of GISTIC and BISCUT loci for COSMIC and Davoli driver genes.

**Supplementary Figure 15.**
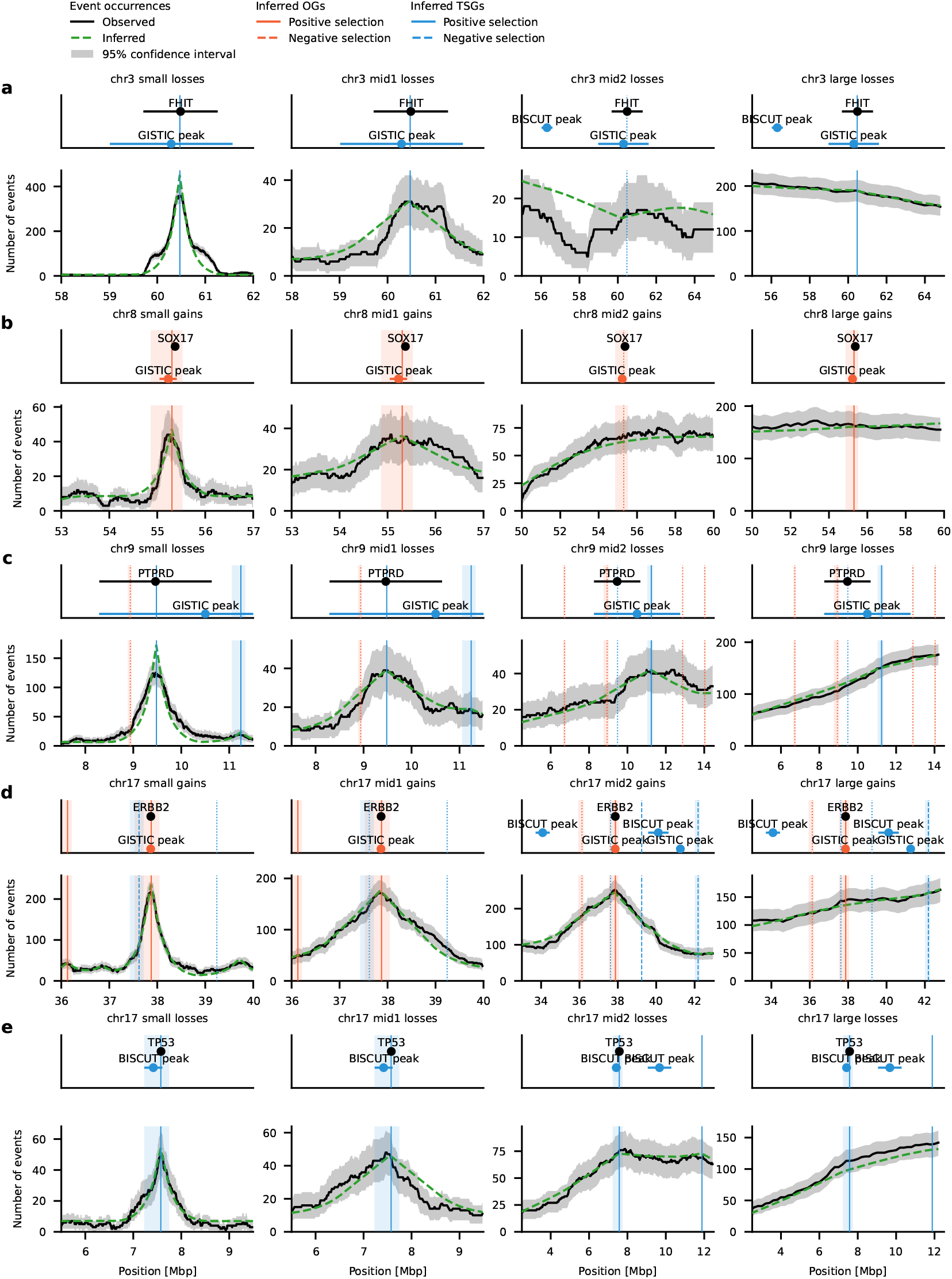
Full length-scale profiles for SPICE loci also identified by GISTIC or BISCUT. Event occurrence profiles across all four event-length classes (small, mid1, mid2, large) for selected loci previously reported by GISTIC or BISCUT: FHIT (chr3), SOX17 (chr8), PTPRD (chr9), ERBB2 (chr17), and TP53 (chr17).

**Supplementary Figure 16.**
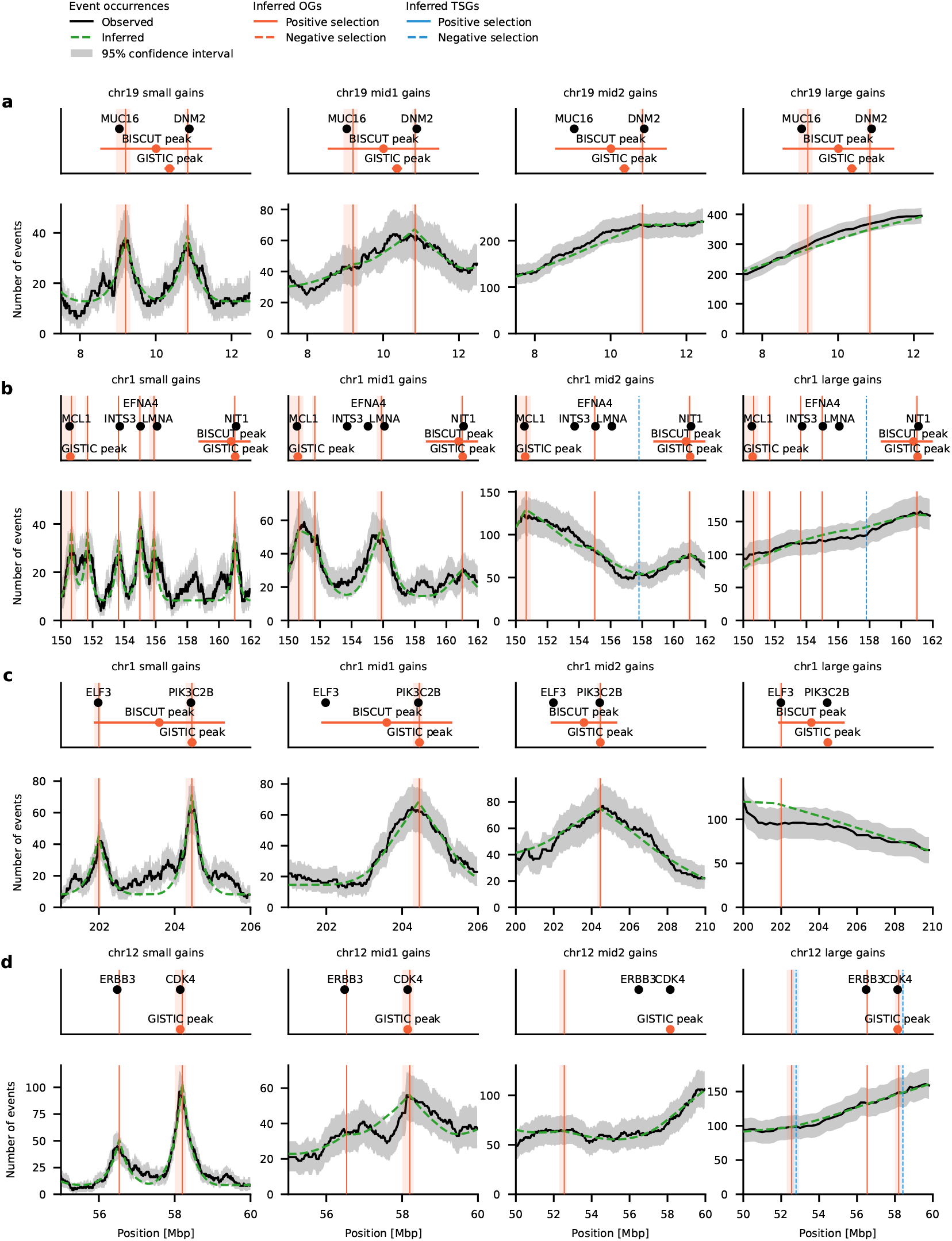
Multi-scale resolution of overlapping selection signals enables finer locus separation by SPICE. SPICE distinguishes closely spaced driver loci that are merged or misplaced in previous methods through its multi-length-scale modeling approach. a, On chromosome 19 (10Mbp), both GISTIC and BISCUT report a single locus; however, SPICE resolves two distinct oncogene-like loci corresponding to MUC16 and DNM2, which converge into one at larger scales. **b**, On chromosome 1 (150–162Mbp), SPICE separates seven individual loci from two broad GISTIC regions, six of which contain known oncogenes, including IFI6 and NT5C. **c–d**, Two more examples on chromosome 1 and 12 where the small length scale uncovers loci of selection that are otherwise hidden.

**Supplementary Figure 17.**
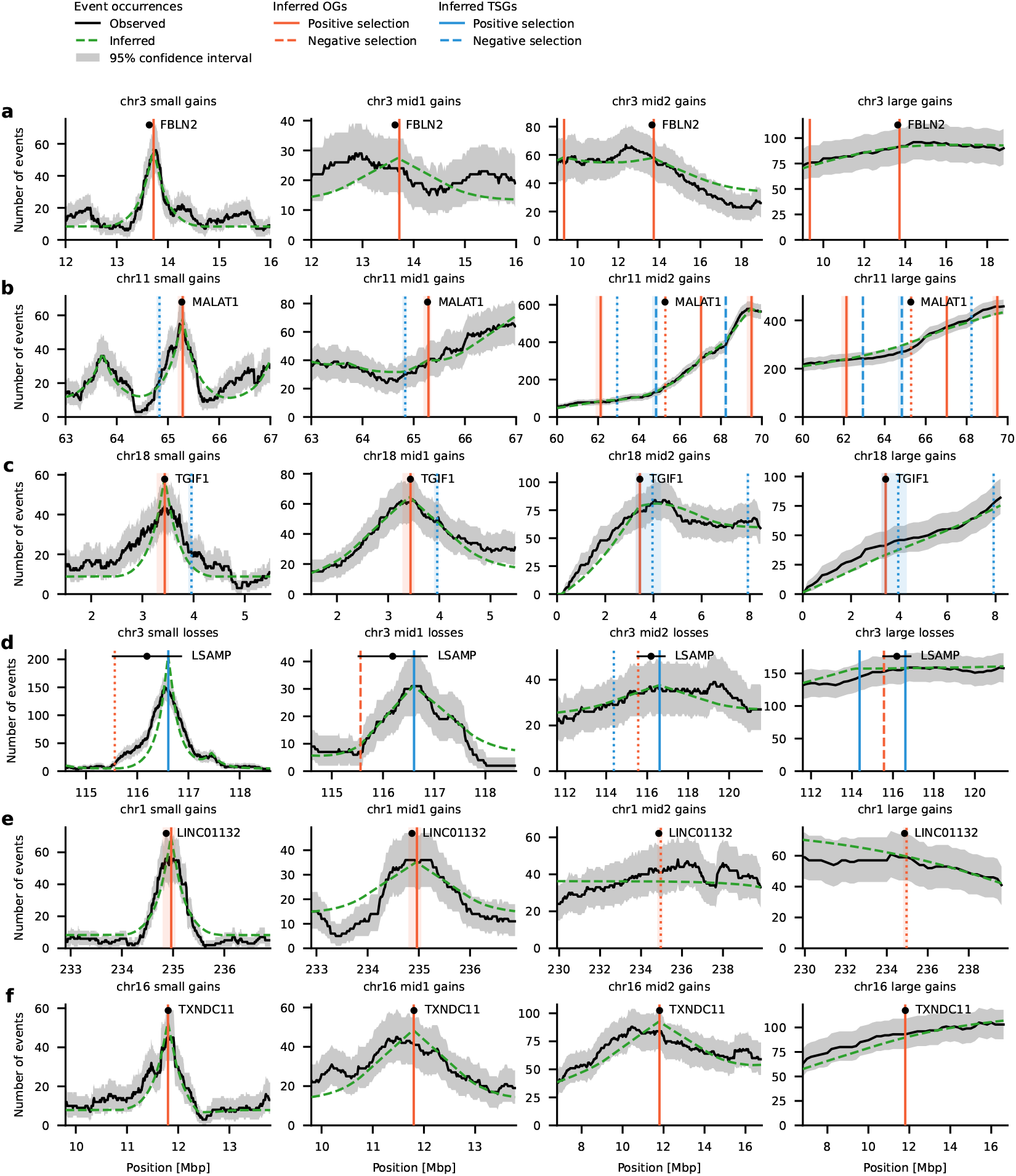
Full length-scale profiles for loci highlighted in Figure 3d. Expanded view of SPICE-inferred loci across all four event-length classes (small, mid1, mid2, large) for the examples shown in the main figure: RBLN2 (chr3), MALAT1 (chr11), TS1F1 (chr18), LSAMP (chr3), LINC01132 (chr1), and TXNDC11 (chr16).

**Supplementary Figure 18.**
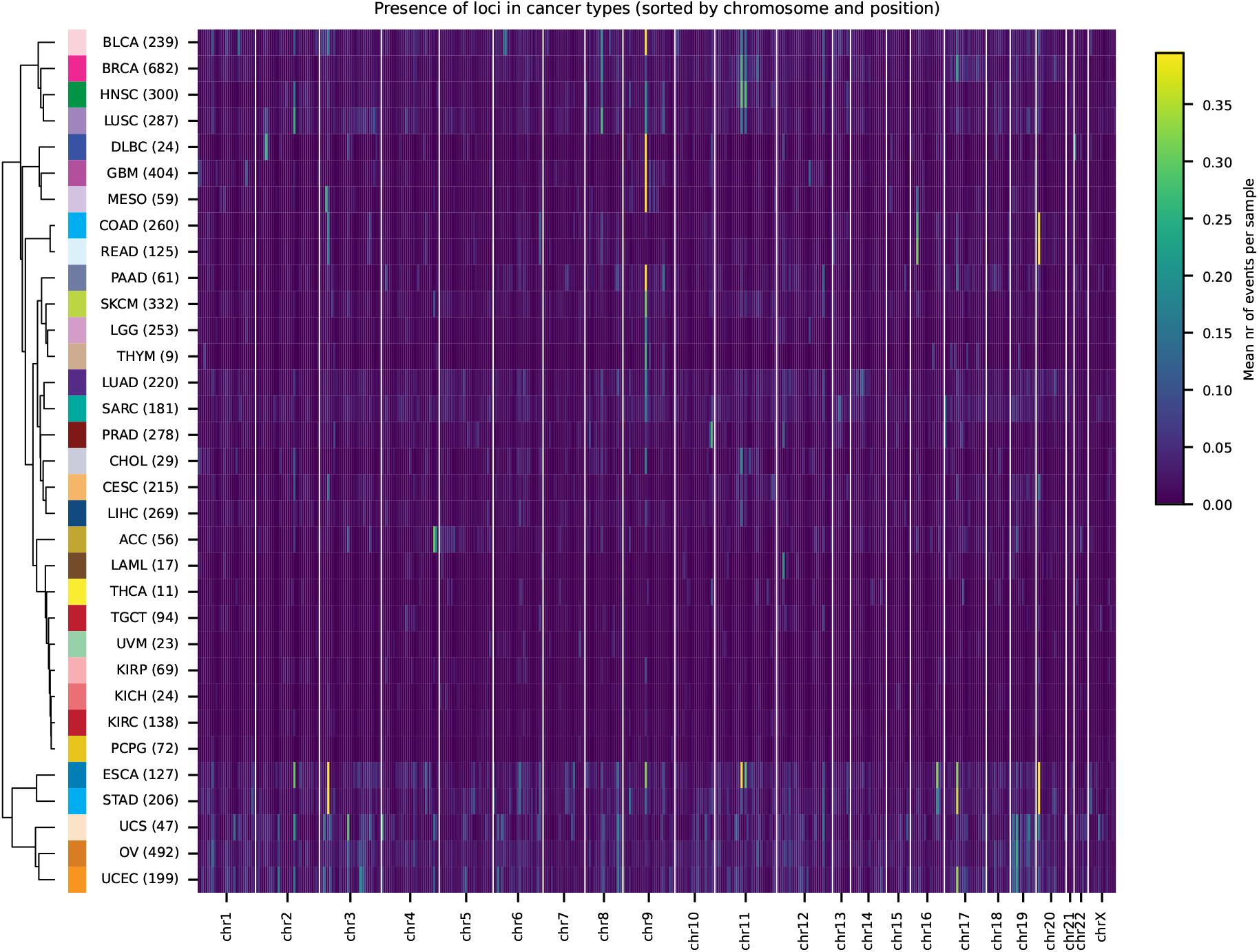
Cancer type–specific patterns of locus-level selection. Heatmap showing the mean number of events per sample for all loci across 33 TCGA cancer types, sorted by chromosome and genomic position. Each row represents a cancer type, and colour intensity indicates the average event frequency at each locus. Cancer types are hierarchically clustered based on similarity of their locus-level profiles. Sample numbers per cohort are shown in parentheses.

**Supplementary Figure 19.**
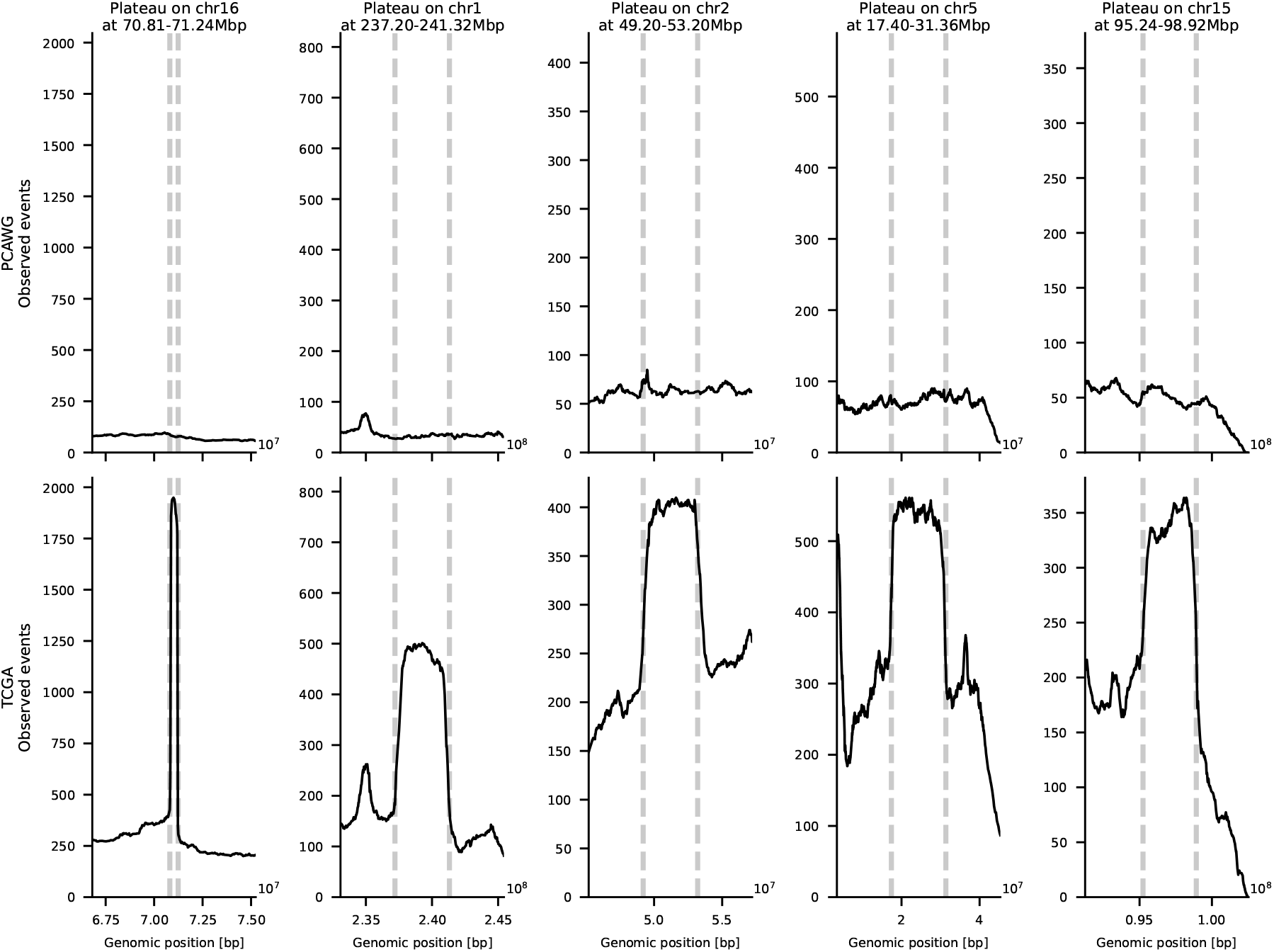
Examples of recurrent technical plateaus in copy-number event signals. Examples of rectangular, step-like artefacts (“plateaus”) detected in aggregated copy-number event profiles. Shown are five representative regions from PCAWG (top) and TCGA (bottom) datasets. In TCGA, plateaus appear as sharp-edged, flat-topped enrichments with near-identical breakpoints across many samples, while these features are absent in the high-quality PCAWG data. Dashed grey lines mark plateau boundaries. Note that the observed events are from the subset of patients that are present in TCGA and PCAWG and the different events counts only correspond to differences in data processing.

