## Supplementary Methods for "Deciphering selection patterns of somatic copy-number events"

### Appendix for the publication “Deciphering selection patterns of somatic copy-number events”

[1]Institute for Computational Cancer Biology (ICCB), Center for Integrated Oncology (CIO), Cancer Research Center Cologne Essen (CCCE), Faculty of Medicine and University Hospital Cologne, University of Cologne, Cologne, Germany.; [2]BIFOLD — Berlin Institute for the Foundations of Learning and Data, Berlin, Germany.; [3]Department of Electrical Engineering and Computer Science, Technische Universität Berlin, Berlin, Germany.; [4]Cancer Research UK Cambridge Institute, University of Cambridge, Cambridge, UK.; [5]Department of Genetics, The University of Texas MD Anderson Cancer Center, Houston, TX, USA.; [6]The Francis Crick Institute, London, UK. ; [7]Department of Genomic Medicine, The University of Texas MD Anderson Cancer Center, Houston, TX, USA.

#### Appendix: Proof of the event reconstruction

The aim of SPICE is to infer a set of copy-number alterations (CNAs) whose combined effect yields a given copy-number (CN) profile.

Any finite set of alterations that induces (reconstructs) the observed CN profile is called a *reconstruction*. In our framework, each alteration is represented as an interval on the genome and a sign $A = \mathbb{N}_1 \times \mathbb{N}_1 \times \{+, -\}$. A gain alteration $(s, e, +)$ increases the copy number by $1$ on the interval $[s, e)$, while a loss alteration $(s, e, -)$ decreases the copy number by $1$ on $[s, e)$. A reconstruction $R \subset A$ is then a finite set of such alterations.

Let $B = (b_1, \dots, b_m)$ be a strictly increasing sequence of breakpoint obtained by sorting the set $\{b \mid \exists (s, e, \pm) \in R : b = s \vee b = e\}$ of start and end positions of all alterations.

The CN profile is then a sequence of copy number values, obtained as a sum of gains and losses that overlap each breakpoint in $B$, i.e. $P = (1, p_1, \dots, p_{|B|})$, where

$$p_i = 1 + \left|\{(s, e, +) \in R : s \le b_i < e\}\right| - \left|\{(s, e, -) \in R : s \le b_i < e\}\right|.$$

Note that $p_{|B|} = 1$ always, because there is no alteration that ends strictly after the last breakpoint. The leading and trailing $1$s represent neutral segments from the start and to the end of the chromosome respectively. If alterations overlap with the ends of a chromosome, the leading or trailing $1$s can be thought of as a zero-width “padding” that ensures that each CN profile starts and ends at $1$. In particular a chromosome with no alterations has profile $P = (1)$ and a chromosome with a single gain will have a profile $P = (1, 2, 1)$ independently of whether it is a full-chromosome, telomere-bound, or internal gain.

##### From copy-number profiles to a unit-step sequence

Let the (padded) CN profile be written as $P = (p_0, \dots, p_{|B|})$, with $p_0 = p_{|B|} = 1$. We first convert $P$ into the sequence of differences

$$\Delta P = (\Delta p_1, \dots, \Delta p_{|B|}), \qquad \Delta p_i := p_i - p_{i-1}.$$

This transformation is bijective: given $P$ we obtain $\Delta P$ by differences, and conversely we can reconstruct $P$ by a cumulative sum starting at $p_0 = 1$. Note that by definition, the $\Delta p_i$ is the copy number change at the breakpoint $b_i$. A reference example is given in

Fig. 1.

If several alterations share a breakpoint, the absolute difference $|\Delta p_i|$ can exceed 1. We decompose such differences into unit changes using a step function $\mathbf{s} : \mathbb{Z} \to \{1, -1\}^*$. We define

$$\mathbf{s}(\Delta p) = \begin{cases} (-1)_{|\Delta p|} & \text{if } \Delta p < 0, \\ (1)_{\Delta p} & \text{if } \Delta p > 0, \end{cases}$$

and construct the unit-step sequence $\delta P = \big(\mathbf{s}(\Delta p_1), \dots, \mathbf{s}(\Delta p_{|B|})\big)$.

This is an alternative representation of the same information. We can convert $\delta P$ back to $\Delta P$ by grouping consecutive unit steps that originated from the same breakpoint and summing them. To keep track of the indices we also use a mapping function $\mathbf{m} : \{1, \dots, |\delta P|\} \to \{1, \dots, |\Delta P|\}$ that maps each index in $\delta P$ to its corresponding index in $\Delta P$. We can then convert $\delta P$ back to $\Delta P$ by summing over the blocks defined by $\mathbf{m}$: $\Delta p_i = \sum_{1 \le k \le |\delta P| : \mathbf{m}(k) = i} \delta p_k$.

#### The bipartite graph

We now encode the unit-step sequence $\delta P$ as a bipartite graph, i.e. a graph whose vertices can be divided into two disjoint sets such that no two graph vertices within the same set are connected. For every index $k$ with $\delta p_k = +1$ we create a vertex $u_k \in U$, and for every index $k$ with $\delta p_k = -1$ we create a vertex $v_k \in V$:

$$U = \{u_k : \delta p_k = +1\}, \qquad V = \{v_k : \delta p_k = -1\}.$$

The final graph $G = (U, V, E)$ is a complete bipartite graph, i.e. $E = U \times V$.

Because $p_0 = p_n$, and each alteration creates exactly one $+1$ and one $-1$, the total number of copy-number increments equals the total number of decrements, and therefore the number of unit gains equals the number of unit losses: $|U| = |V|$.

Each edge $(u_i, v_j) \in E$ will represent a single CNA. If the $+1$ occurs before the $-1$, the alteration is a gain; otherwise, it is a loss. Formally, $(u_i, v_j) \in E \wedge i < j \implies (b_{\mathbf{m}(i)}, b_{\mathbf{m}(j)}, +) \in R$ and $(u_i, v_j) \in E \wedge j < i \implies (b_{\mathbf{m}(j)}, b_{\mathbf{m}(i)}, -) \in R$.

#### Reconstructions of $\delta P$

Each unit increase in $\delta P$ must be paired with a unit decrease to form an alteration. We therefore formalize reconstructions at the level of the unit-step sequence.

**Definition (Reconstruction of $\delta P$).** A finite set of alterations is a *reconstruction of* $\delta P$ if

1. each alteration corresponds to exactly one $+1$ and one $-1$ in $\delta P$ (i.e. it contributes $+1$ at a single index and $-1$ at a single index, and $0$ elsewhere), and
2. for every index $k$ with $\delta p_k = +1$ there is exactly one alteration that contributes $+1$ at position $k$, and for every index $k$ with $\delta p_k = -1$ there is exactly one alteration that contributes $-1$ at position $k$.

Equivalently, the unit increases and decreases in $\delta P$ are paired up bijectively by the alterations.

We say that a set of edges $M \subseteq E$ *encodes* a reconstruction of $\delta P$ if the alterations obtained from $M$ by the construction below form a reconstruction in this sense.

##### Example

Consider an example of one p-arm, and one centromere-bound gain on chromosome 1, i.e. $R = \{(1, 125{\cdot}10^6, +), (50{\cdot}10^6, 125{\cdot}10^6, +)\}$. The $R$ induces the strictly ordered set of breakpoints $B = (1, 50 \cdot 10^6, 125 \cdot 10^6)$, and the profile $P = (1, 2, 3, 1)$,

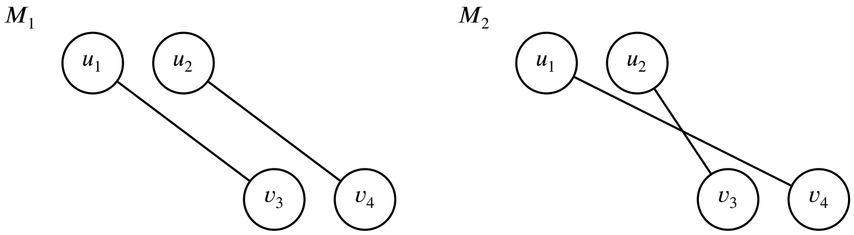


**Appendix Figure 1.** The two perfect matchings for the profile $P = (1, 2, 3, 1)$. Although the pairings differ, $v_3$ and $v_4$ correspond to the same breakpoint ($b_{\mathbf{m}(3)} = b_{\mathbf{m}(4)}$), so both matchings induce the same set of alterations.

---

###### Step 1: Difference sequence.

Computing the differences $\Delta p_i = p_i - p_{i-1}$ yields $\Delta P = (+1, +1, -2)$.

###### Step 2: Unit-step sequence.

The entry $\Delta p_3 = -2$ is decomposed into two unit decreases $\delta P = (+1, +1, -1, -1)$. The mapping from the sequence of unit steps to the sequence of defferences gives $\mathbf{m}(1) = 1$, $\mathbf{m}(2) = 2$, and $\mathbf{m}(3) = \mathbf{m}(4) = 3$.

###### Step 3: Vertex sets.

From $\delta P$ we obtain

$$U = \{u_1, u_2\} \quad \text{(positions with } +1), \qquad V = \{v_3, v_4\} \quad \text{(positions with } -1).$$

###### Step 4: Bipartite graph and matchings.

The complete bipartite graph $G = (U, V, E)$ has edge set $E = \{(u_1, v_3), (u_1, v_4), (u_2, v_3), (u_2, v_4)\}$. There are exactly two perfect matchings, shown in Fig. 1.

##### Perfect matchings encode reconstructions

We now show that every perfect matching in $G$ encodes a reconstruction of $\delta P$ and, conversely, that every reconstruction of $\delta P$ defines a perfect matching in $G$.

**Lemma 0.1.** *If $M$ is a perfect matching in $(U, V, E)$ then $M$ encodes a reconstruction of $\delta P$.*

*Proof.* Because $M$ is a perfect matching, every vertex in $U \cup V$ has degree 1 in $M$. For each edge $e = (u_i, v_j) \in M$ we construct a single alteration as follows:

1. If $i < j$ we create a gain alteration that increases the copy number on the genomic interval from $b_{\mathbf{m}(i)}$ to $b_{\mathbf{m}(j)}$. At the level of the unit-step sequence, this alteration contributes $+1$ at index $i$ and $-1$ at index $j$ of $\delta P$.
2. If $j < i$ we create a loss alteration that decreases the copy number on the genomic interval from $b_{\mathbf{m}(j)}$ to $b_{\mathbf{m}(i)}$. At the level of $\delta P$, this alteration contributes $-1$ at index $j$ and $+1$ at index $i$.

In either case, the alteration associated with $(u_i, v_j)$ contributes exactly one $+1$ and one $-1$ to the unit step sequence and $0$ at all other indices.

Now fix any index $k$ with $\delta p_k = +1$. Because $M$ is a perfect matching, there is a unique edge $(u_k, v) \in M$ incident to $u_k$, and the corresponding alteration is the only alteration that contributes $+1$ at position $k$.

Similarly, for any index $k$ with $\delta p_k = -1$, there is a unique edge $(u, v_k) \in M$ incident to $v_k$. The corresponding alteration is the only alteration that contributes $-1$ at position $k$.

Thus the set of alterations obtained from $M$ uses each unit increase and each unit decrease in $\delta P$ exactly once and yields the desired unit step sequence. Hence $M$ encodes a reconstruction of $\delta P$. □

In the following we say that $M \subseteq E$ *encodes a reconstruction of* $\delta P$ if the alterations constructed from $M$ satisfy the two conditions in the definition above: each alteration contributes exactly one $+1$ and one $-1$, and each unit step in $\delta P$ is used by exactly one alteration.

**Lemma 0.2.** *Let $M \subseteq E$ be a set of edges encoding a reconstruction of $\delta P$. Then $M$ is a perfect matching in $(U, V, E)$.*

*Proof.* Consider any index $k$ with $\delta p_k = +1$. Since $M$ encodes a reconstruction of $\delta P$, exactly one alteration contributes $+1$ at position $k$. This alteration must arise from a unique edge in $M$ incident to $u_k$. Hence $u_k$ has degree $1$ in $M$.

Similarly, for any index $k$ with $\delta p_k = -1$ there is exactly one alteration that contributes $-1$ at position $k$, so there is a unique edge in $M$ incident to $v_k$. Thus $v_k$ also has degree $1$ in $M$.

We conclude that every vertex in $U \cup V$ has degree exactly $1$ in $M$. Therefore $M$ is a perfect matching in $(U, V, E)$. □

#### Minimal number of alterations

We next show that any reconstruction of $\delta P$ must contain at least $|U|$ alterations.

**Lemma 0.3.** *Any reconstruction of $\delta P$ requires at least $|U|$ alterations.*

*Proof.* By construction, the sequence $\delta P$ contains

$$|U| = \left|\{k : \delta p_k = +1\}\right|$$

positions at which the copy number increases by $1$, and

$$|V| = \left|\{k : \delta p_k = -1\}\right| = |U|$$

positions at which it decreases by $1$.

In any reconstruction of $\delta P$, each alteration contributes exactly one $+1$ and one $-1$ to the unit step sequence. Hence the total number of $+1$ contributions made by all alterations equals the number of alterations in the reconstruction.

On the other hand, to obtain the desired sequence $\delta P$ we must realize all $|U|$ positions with $\delta p_k = +1$. Each such position is accounted for by exactly one alteration. Therefore the number of alterations is at least $|U|$. More alterations are can always be created by either splitting existing alterations into smaller ones or by adding pairs of symmetric gains and losses.

□

#### One-to-one correspondence and upper bound

Combining the previous lemmas yields a bijection between perfect matchings in $G$ and minimal reconstructions of the unit-step sequence. After mapping the nodes to breakpoints, some solutions may create the same set of alterations. An example of two solutions that create the same reconstruction is shown in Fig. 1.

In practice, we treat different chromosomes independently and consider reconstructions as unordered sets of CNAs. Thus permuting alterations across chromosomes does not yield a distinct reconstruction.

**Theorem 0.4.** *There is a one-to-one correspondence between perfect matchings in $(U, V, E)$ and minimal reconstructions of the unit-step sequence $\delta P$ (and hence of $P$ at the level of unit steps).*

*Proof.* Lemma 0.1 shows that every perfect matching in $G$ encodes a reconstruction of $\delta P$, and hence (by inverting the transformations above) a reconstruction of $P$. Conversely, Lemma 0.2 shows that every reconstruction of $\delta P$ corresponds to a perfect matching. By Lemma 0.3, any reconstruction must contain at least $|U|$ alterations, so perfect matchings correspond precisely to minimal reconstructions at the unit-step level. □

**Corollary 0.5.** *There are at most $|U|!$ minimal reconstructions of $\delta P$.*

This upper bound is obtained by observing that a perfect matching in $(U, V, E)$ defines a bijection between $U$ and $V$. There are at most $|U|!$ such bijections, and therefore at most $|U|!$ perfect matchings and at most $|U|!$ minimal reconstructions of the unit-step sequence.

#### Loss of segments

In some cases, segments can be completely lost, i.e. the copy number becomes $0$ and can no longer be regained. However, in our framework we do not model concatenation of neighbouring segments, as would occur in internal chromosomal deletions. This can create spurious alterations: when the two neighbours of a lost segment are gained together, two alterations are created instead of one.

We avoid this in SPICE by allowing gains from $0$ in cases where two neighbours of a lost segment are gained simultaneously. We assume that whenever both neighbours of a lost segment are gained, they are gained together. For each such case, we adjust the profile by adding a segment that connects the two neighbours. Formally, we create $\bar{P} = (1, \bar{p}_1, \ldots, \bar{p}_{n-1}, 1)$ from $P = (1, p_1, \ldots, p_{n-1}, 1)$ as follows:

$$\bar{p}_i = \begin{cases} \min(p_{i-1}, p_{i+1}) & \text{if } p_i = 0, \\ p_i & \text{otherwise.} \end{cases}$$

We then reconstruct $\bar{P}$ using the graph-based procedure above. In this way we avoid spurious alterations around lost segments. However, we must then ensure that there exists an ordering of alterations such that segments with copy number $0$ are indeed lost at least once (reach CN 0) and all remaining segments are never lost (never reach CN 0). This condition is enforced in a post-processing step using a constraint solver, as detailed in the main Methods.

#### Whole-genome duplication

Our method does not explicitly detect whole-genome duplication (WGD) alterations, where pre-WGD gains will appear as two overlapping gains post-WGD, creating 2 edges in the graph representation. Under the minimum evolution assumption, two overlapping gains in a sample with WGD are assumed to be a pre-WGD gain, since this results in fewer total alterations.

We treat samples labelled as WGD in the dataset separately from non-WGD samples. Since the neutral CN after WGD is $2$ per allele, we set the baseline copy number to $2$, i.e. we consider $P2 = (2, p2_1, \ldots, p2_{|B|})$ and define

$$p2_i = 2 + \left|\{(s, e, +) \in R : s \le b_i < e\}\right| - \left|\{(s, e, -) \in R : s \le b_i < e\}\right|.$$

A pre-WGD gain induces two consecutive increments and two consecutive decrements in $\delta P2$, in an order depending on whether it is a gain or loss. In the reconstruction we distinguish between pre- and post-WGD alterations by partitioning the edges of a perfect matching $M$ into two disjoint sets $R_{\text{pre}}$ and $R_{\text{post}}$, with $R_{\text{pre}} \cup R_{\text{post}} = M$ and $R_{\text{pre}} \cap R_{\text{post}} = \emptyset$. We require that

$$(u_i, v_\ell) \in R_{\text{pre}} \implies (u_{i+1}, v_{\ell+1}) \in R_{\text{pre}} \text{ or } (u_{i-1}, v_{\ell-1}) \in R_{\text{pre}},$$

and that $|R_{\text{pre}}|$ is even. Thus $R_{\text{pre}}$ is a union of disjoint pairs of neighbouring edges representing duplicated pre-WGD alterations.

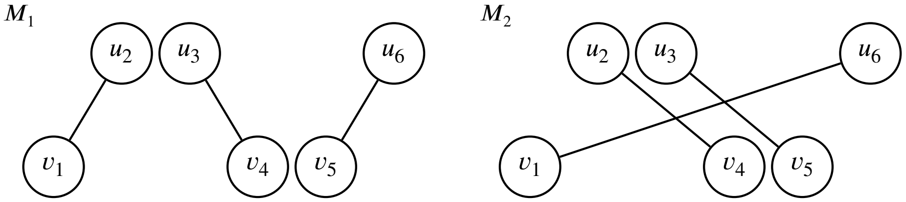


**Appendix Figure 2.** Two perfect matchings in a bipartite graph representing the profile $P = (2, 1, 3, 1, 2)$ and the unit-step sequence $\delta P = (-1, +1, +1, -1, -1, +1)$. The matching $M_1$ represents three post-WGD alterations. The matching $M_2$ represents one pre-WGD and one post-WGD alteration.

---

As illustrated in Fig. 2, two different perfect matchings can represent the same set of genomic alterations, depending on whether the alterations are classified as pre- or post-WGD. We estimate the total number of alterations using MEDICC with the `x2` option enabled (see main Methods). For a profile with $m$ estimated alterations, we only retain reconstructions satisfying

$$\frac{|R_{\text{pre}}|}{2} + |R_{\text{post}}| = m.$$

In practice, when constructing a matching, admissible pairs of neighbouring edges are first assigned to $R_{\text{pre}}$ (subject to the pairing rule above) until the constraint is met; the remaining edges are assigned to $R_{\text{post}}$.

In a case of simultaneous LoH and WGD, we apply the constraint solving separately for the pre- and post-WGD alterations.

#### Shared breakpoints in WGD samples

Finally, we consider shared telomere breakpoints in WGD samples. A breakpoint may be explained either by a post-WGD gain followed by a loss or by a pre-WGD gain followed by a post-WGD loss. Both possibilities correspond to two alterations, but the first is represented by two edges in the alteration graph, whereas the second is represented by three. To capture both scenarios, we construct an additional graph $G' = (U', V', E')$ that contains an extra pair of vertices.

Intuitively, in the original graph $G$, the gain-and-loss breakpoint can be seen as a ∧-shape or ∨-shape, depending on the order of gain and loss. We place the additional pair of vertices in $G'$ that represent a zero-width alteration sharing the same breakpoint, as illustrated in Fig. 3.

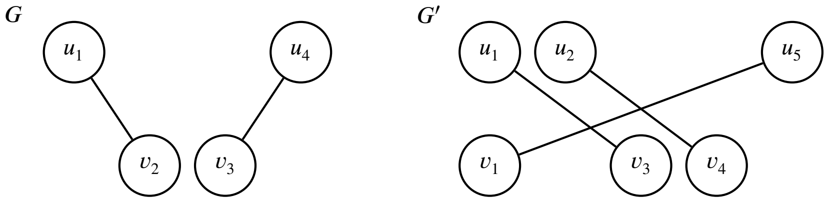


**Appendix Figure 3.** Shared telomere breakpoints in WGD samples. Left: Graph $G$ with vertices $u_1, v_2, v_3, u_4$ and $b_{\mathbf{m}(2)} = b_{\mathbf{m}(3)}$, representing a post-WGD gain and loss. Right: Graph $G'$ with additional vertices for a zero-size alteration representing a pre-WGD gain and post-WGD loss scenario.

Formally, for a set $u_i, v_{i+1}, v_{i+2}, u_{i+3}$ in $G$ such that $\mathbf{m}(v_{i+1}) = \mathbf{m}(v_{i+2})$ (representing gain-and-loss), we first increment all indices after $i$ and then insert $u'_i, v'_i$ such that:

$$U' = \{u'_k : u_k \in U, k < i\} \cup \{u'_{k+1} : u_k \in U, k \geq i\} \cup \{u'_i\},$$

$$V' = \{v'_k : v_k \in V, k < i\} \cup \{v'_{k+1} : v_k \in V, k \geq i\} \cup \{v'_i\}.$$

The process is analogous for a set $v_i, u_{i+1}, u_{i+2}, v_{i+3}$ in $G$ such that $\mathbf{m}(u_{i+1}) = \mathbf{m}(u_{i+2})$ (representing a loss-and-gain), however there we insert the new edges at the position $i + 4$ instead of $i$ and increment all previous indices after $i + 3$.

In case there are multiple possible shared breakpoints, we create all possible combinations of graphs with added vertices, e.g. for two breakpoints $b_i, b_j$ we create graphs $G'_{b_i}$, $G'_{b_j}$ and $G'_{b_i,b_j}$ in addition to the original graph $G$.
